# Chemotaxing *E. coli* do not count single molecules

**DOI:** 10.1101/2024.07.09.602750

**Authors:** Henry H. Mattingly, Keita Kamino, Jude Ong, Rafaela Kottou, Thierry Emonet, Benjamin B. Machta

## Abstract

Organisms use specialized sensors to measure their environments, but the fundamental principles that determine their accuracy remain largely unknown. In *Escherichia coli* chemotaxis, we previously found that gradient-climbing speed is bounded by the amount of information that cells acquire from their environment, and that *E. coli* operate near this bound. However, it remains unclear what prevents them from acquiring more information. Past work argued that *E. coli*’s chemosensing is limited by the physics of molecules stochastically arriving at cells’ receptors, without direct evidence. Here, we show instead that *E. coli* are far from this physical limit. To show this, we develop a theoretical approach that uses information rates to quantify how accurately behaviorally-relevant signals can be estimated from available observations: molecule arrivals for the physical limit; chemotaxis signaling activity for *E. coli*. Measuring these information rates in single-cell experiments across multiple background concentrations, we find that *E. coli* encode two orders of magnitude less information than the physical limit. Thus, *E. coli* chemosensing is limited by internal noise in signal processing rather than the physics of molecule diffusion, motivating investigation of what specific physical and biological constraints shaped the evolution of this prototypical sensory system.

## Introduction

Evolution selects function, and therefore living systems are shaped by complex fitness objectives and constraints. This has motivated the use of normative theories, subject only to constraints of physics, to derive fundamental limits on function and to rationalize the design of biological systems (1–17). This approach has been especially successful in the context of information processing, a hallmark of living systems where theories of optimal estimation can be brought to bear (18,19). However, biology needs to implement information processing and other functions using non-ideal components, in the confines of a body, and with limited resources, which introduce additional system-specific constraints (20–27). Determining what bounds or constraints meaningfully limit information processing in a particular biological system would shed light on the forces that have shaped its evolution, and inform our understanding of biological information processing more broadly.

*Escherichia coli* chemotaxis is an ideal system for studying the limits on biological information processing (28–30). *E. coli* climb chemical gradients by alternating between straight-swimming runs and randomly-reorienting tumbles (31). As they swim, they measure the time-dependent concentration of attractant along their trajectory, *c(t)*, using transmembrane receptors, encode these measurements into the activity of intracellular, receptor-associated CheA kinase activity, *a(t)*, and act on these measurements to decide when to tumble (Fig. 1). Importantly, chemotaxis provides a fitness advantage, even above undirected motility, in structured chemical environments (32).

**Figure 1.**
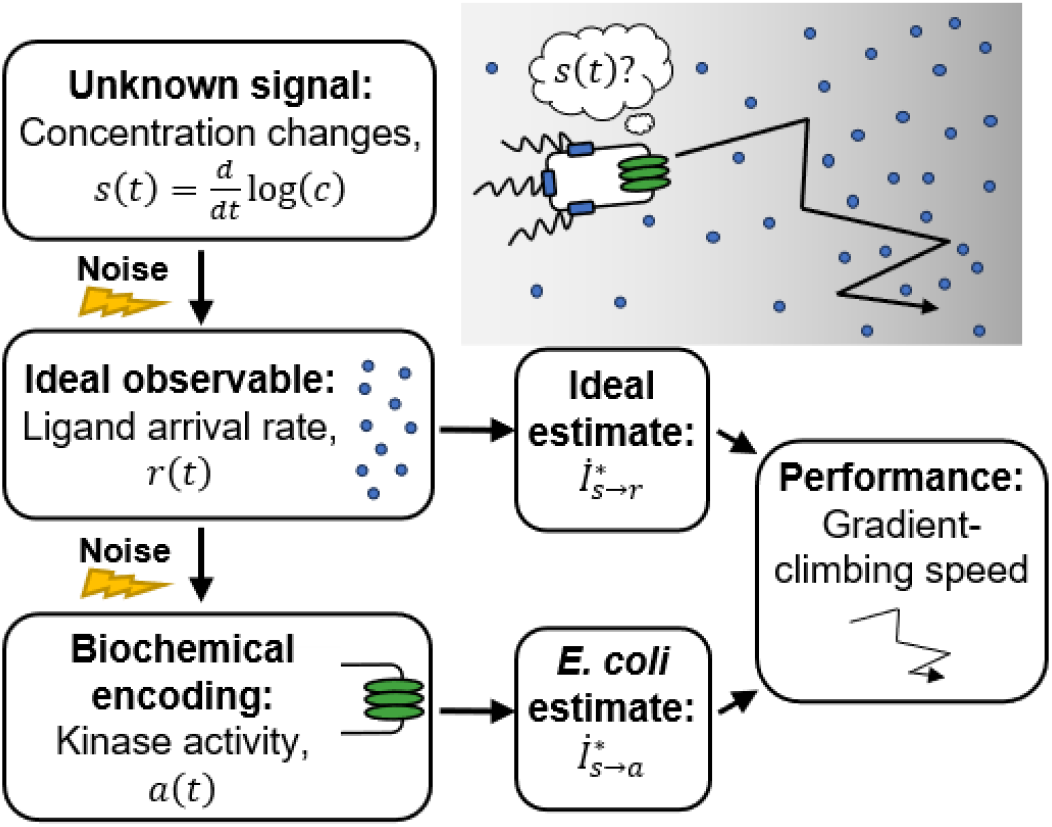
Is *E. coli*’s sensing accuracy set by physical limits or internal constraints? To climb chemical gradients, *E. coli* need to accurately estimate an unknown signal: the rate of change of attractant ligand concentration,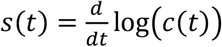 (17). The first physically-observable quantity is the stochastic rate at which ligand molecules arrive at the cell’s receptors, *r(t)*(4). Thus, the physical limit on chemosensing, and in turn gradient-climbing speed, is set by how accurately *s(t)* can be estimated from the time series of past *r(t)*, quantified by an information rate,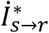. *E. coli* respond to ligand arrivals with changes in the activity of intracellular CheA kinases, *a(t)*. The accuracy with which the signal can be estimated from kinase activity is quantified by another information rate, 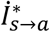. Since kinase activity is stochastic, *E. coli*’s sensing accuracy and gradient climbing speed must be below the physical limit, but how much less?

*E. coli* must acquire information about their chemical environment in order to climb gradients. Recently, we asked how fast an ideal bacterium can climb a gradient with the information it gets, and how *E. coli* compare to this theoretical performance bound (17). We found that although typical *E. coli* cells get very little information about chemical signals—about 0.01 bits/s in a centimeter-long gradient—they climb gradients at speeds near the theoretical maximum with the information they get. Thus, information is functionally important for chemotaxis.

This raises the question: why don’t *E. coli* get more information, and thus climb gradients faster? One possibility is that they are limited by fundamental physics. The first physically-measurable quantity is the rate of ligand molecule arrivals at the cell’s receptors by diffusion, *r(t)*(Fig. 1). In a classic paper (4), Berg and Purcell demonstrated that the stochasticity of this arrival rate limits the accuracy of any estimate of chemical concentration, *c(t)*, inspiring an entire field of biophysics (20,21,33–47). They and others further argued that bacteria approach this physical limit, a widely-held understanding in the field. However, no direct comparison between bacterial chemosensing and physical limits has been made because it has remained unclear how to quantify a real cell’s uncertainty about external signals. This leaves open the alternative possibility that *E. coli*’s sensory information might be limited by system-specific, internal constraints.

Directly answering whether physical limits or internal constraints prevent chemotaxing *E. coli* from acquiring more information faces several general challenges. First, not all environmental signals are useful for function. For chemotaxis in shallow gradients, we recently showed that the time derivative of (log) concentration 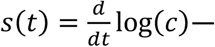 rather than concentration, *c(t)*, itself—is the “behaviorally-relevant” signal (17). Second, cells do not need to represent their estimates of relevant signals in a straightforward way. In chemotaxis, CheA kinase activity depends on external signals, but *a(t)* is not necessarily the cell’s estimate of *s(t)*, and the variation in *a(t)* is not the cell’s uncertainty about *s(t)*. Signals are instead encoded in the dynamics of the cell’s intermediate variables and decoded by downstream processing. Third, probing cells’ encodings of time-varying signals requires dynamic experimental measurements of both the environment and responses in single cells, which was recently made possible by single-cell FRET (17,48–52).

Here, we address these challenges and determine whether *E. coli* chemosensing approaches the physical limits. To frame this question in an experimentally-testable way, we ask how accurately the signal *s(t)* can be inferred from molecule arrivals, which sets the physical limit, compared to how accurately *s(t)* can be inferred from the dynamics of kinase activity, the quantity accessible to the cell. Sensing accuracy in each case takes the form of an information rate (Fig. 1). Then, we quantify these information rates using single-cell FRET measurements in multiple background concentrations. We find, surprisingly, that a typical *E. coli* cell gets orders of magnitude less information than the physical limit—estimates of signal made from kinase activity are far less accurate than those made from molecule arrival rate. This is because *E. coli*’s signal transduction noise far exceeds molecule arrival noise, and we conclude that information processing during *E. coli* chemotaxis is *internally-limited*. We predict that the functional consequence is that *E. coli* climb gradients much slower than the physical limits on chemosensing allow, and support this with simulations. These results raise questions about what specific constraints limit *E. coli*’s chemosensing, and more broadly motivate consideration of the physical and biological constraints on information processing.

### Physical limit on behaviorally-relevant information due to stochastic molecule arrivals

To climb chemical gradients, *E. coli* must encode information about the time derivative of concentration, *s(t)*, to be read out by the motors (17) (SI section “Drift speed and information rate”). The first quantity that is observable to the cell and informative of *s(t)* is the stochastic arrival rate of ligand molecules at the cell’s receptors, *r(t)*(Fig. 1). An ideal agent would estimate *s(t)* and make navigation decisions based on perfect observations of past particle arrivals {*r*}. The behaviorally-relevant information about signal, *s(t)*, thus acquired from past particle arrivals, *{r*}, is quantified by the following transfer entropy rate (53):

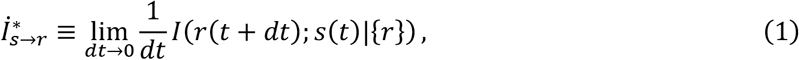

where *I(X; Y*|*Z)* is the mutual information between *X* and *Y*, conditioned on *Z* (54,55). This quantity obeys a data processing inequality (55) in the context of bacterial chemotaxis in shallow gradients, where feedback from behavior onto signals is negligible (56,57) (SI section “Data processing inequality”). Therefore this quantity sets the physical limit on information available in any downstream encoding of the signal, including *E. coli*’s kinase activity.

The form of the physical limit in Eqn. <u>1</u>is unknown. To derive it, we first need a dynamical model for the signal and the particle arrival rate. In static gradients, the signals a cell experiences are determined by their own run-and-tumble motion in the gradient. Accordingly, in a gradient of steepness *g* =*d* log*(c) /dx*, the signal is *s(t)* = *g v*_*x*_*(t)*, where *v*_*x*_ is the cell’s up-gradient velocity. In shallow gradients, where weak signals have small effects on the cell’s run-tumble statistics, we can rigorously approximate *s(t)* as Gaussian with correlation function 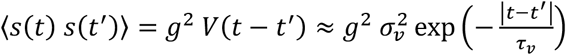, to leading order in *g* (17,22). Here, *V(t)* is the correlation function of *v*_*x*_ in the absence of a gradient; 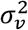 is the variance of *v*_*x*_, which depends on the cell’s swimming speed; and τ*v* is the signal correlation time, which depends on the cell’s mean run duration, the persistence of tumbles, and rotational diffusion (17,58).

Molecule arrival events follow a Poisson process with time-varying rate ⟨*r(t)*⟩ = k_*D*_ *c(t)* = 4 *D* l *c(t)*, where *D* ≈ 800 μm^2^*/*s (59,60) is the diffusivity of the ligand and l is the radius of a circular sensor on the cell’s surface (4,42). We choose l ≈ 60 nm (61) to match the size of the receptor array in *E. coli*’s cell membrane. These give k_*D*_ ≈ 1.2 × 10^5^ s^−1^ μM^−1^, which is comparable to previous estimates (4,62). If many molecules arrive per run, *r*_0_ τ_*v*_ ≫ 1, we can approximate the Poisson process for arrival events with a Gaussian process for the number of molecule arrivals per unit time, 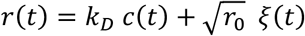. Here, *r*_0_ = k_*D*_ *c*_0_ is the background molecule arrival rate, *c*_0_ is the background concentration, and the noise is ⟨*ξ*(*t*) ξ(*t*′)⟩ = *δ(t* − *t*′). We assume the sensor absorbs every molecule it senses (4), but if it cannot distinguish between new ligand arrivals and rebinding events, the limit is lower by an *O(*1) prefactor (42,43).

We next focused on calculating the behaviorally-relevant information quantity in Eqn. <u>1</u>. Towards this, we discovered that the transfer entropy rate in Eqn. <u>1</u> is equivalent to a predictive information rate (22,23,63– 66) (SI section “Equivalence of transfer entropy and predictive information rates”):

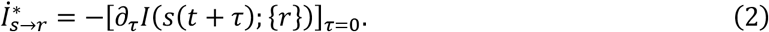

This quantifies how the ability to predict future signals *s(t* + τ) from past particle arrivals degrades as the forecasting interval τ increases, and is evaluated at the current moment, τ = 0. Importantly, this quantity only quantifies the information that is relevant for climbing the gradient. Therefore it is different from the total information encoded by *E. coli*’s signaling pathway about *all* past signals, *{s*}, both relevant (current signal) and irrelevant (signal experienced in the past), that we and others studied previously (17,57,67,68).

Since *s(t)* and *{r*} are approximately Gaussian, the physical limit in Eqn. <u>2</u> only depends on the posterior variance, 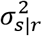, of *s(t)* given past particle arrivals *{r*} (SI Eqn. 23), which can be derived using causal Wiener filtering theory (22,64,69–74) (SI section “Derivation of the physical limit on behaviorally-relevant information for chemotaxis”). We find that the physical limit on behaviorally-relevant information for chemotaxis in shallow gradients is:

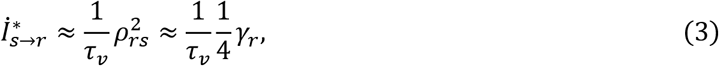

where *ρ*_*rs*_ is the Peason correlation coefficient between the true signal *s(t)* and the optimal estimator of *s(t)* constructed from past molecule arrivals, 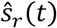. Here, we defined the dimensionless signal-to-noise ratio of molecule arrivals,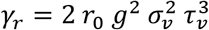. Eqn. <u>3</u> is valid when *γ*_*r*_ ≪ 1, which sets the small-signal regime for 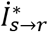. We also provide a full expression for 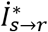 in the SI (SI Eqn. 46), and we validate our expression for 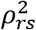 using simulations (SI Fig. S5). Increasing the background *r*_0_, the gradient steepness *g*,or the swimming speed σ_*v*_ increases the signal-to-noise ratio of molecule arrivals. Longer runs, τ_*v*_, also increases 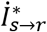 by allowing more time to average out noise. We expect spatial sensing across the cell body to be negligible compared to temporal sensing, as argued by Berg and Purcell (SI section “Comparing temporal and spatial sensing”; see also (75)). The derivation of 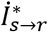 also provides the optimal kernel for constructing 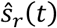, which we discuss in the SI (section “Optimal kernel for estimating signal from particle arrivals”).

### Relevant information encoded in *E. coli*’s CheA kinase activity

In *E. coli*, ligand binding to receptors modulates the activity of the CheA kinases in the receptor-kinase complex. Thus, kinase acivity *a(t)* depends on past signals *s(t)*, but is not necessarily the cell’s representation of them. To compare *E. coli* to the theoretical limit, we next derive 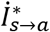, which quantifies how well *s(t)* can be estimated from the dynamics of kinase activity. For this, we need models of kinase responses to ligand molecule arrivals and noise in kinase activity. In shallow gradients, our approach is to use linear, Gaussian theory, which has been validated experimentally (17,48,49) and computationally (68). For a cell with steady-state kinase activity *a*_0_ in background *r*_0_, kinase responses are described by linear response theory (17,76,77) as follows:

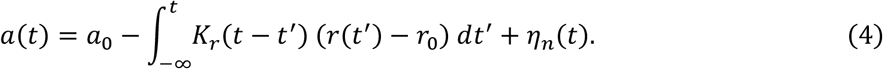

*E. coli* respond to a step increase in attractant concentration with a fast drop in kinase activity, followed by slow adaptation back to the pre-stimulus level (78). We model this phenomenologically with response function 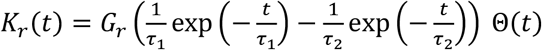, where *G*_*r*_ is the gain of the response to molecule arrival rate *r*, τ_1_ is the fast response time, τ_2_ is the slow adaptation time, and Θ*(t)* is the Heaviside step function. Kinase responses can equivalently be expressed in terms of past signals *s*, with a related kernel *K(t)* that we used previously 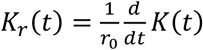.

Noise in kinase activity is driven by a combination of stochastic molecule arrivals and internally-driven fluctuations. Previous single-cell FRET experiments have observed large, slow fluctuations in kinase activity, η_*n*_*(t)*, on a time scale of 10 s (17,48,49,79). These are well-described as Gaussian, with correlation function 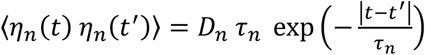. Here,*D*_*n*_ is the diffusivity of internal noise in kinase activity, and τ_*n*_ is its correlation time. In addition, Eqn. <u>4</u> has additive noise arising from responses to molecule arrival noise. To date, it has not been possible to measure kinase fluctuations on time scales shorter than the CheY-CheZ relaxation time (τ_1_), but it cannot go below the level set by responses to molecule arrival noise. Thus, the phenomenological model above agrees with experiments at low frequencies while obeying known physics at high frequencies.

With the relation between transfer entropy and predictive information in Eqn. <u>2,</u> evaluating 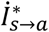 again reduces to deriving the posterior variance, 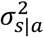, of the signal *s(t)* given past kinase activity *{a*} (SI section “Derivation of the behaviorally-relevant information in kinase activity”). Furthermore, previous measurements (and measurements below) show that τ_1_ ≪ τ_*v*_ (17,80,81) and τ_2_ ≈ τ_*n*_ ≫ τ_1_ (17). Thus, in shallow gradients, we find that the information rate encoded in kinase activity is:

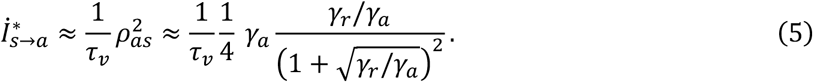

where *ρ*_*as*_ is the Peason correlation coefficient between the true signal *s(t)* and the optimal estimator of *s(t)* constructed from past kinase activity, 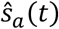. Here, we define the dimensionless kinase activity signal-to-noise ratio 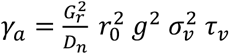. Eqn. <u>5</u> is valid when *γ*_a_ ≪ 1, which sets the small-signal regime for 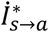. We also provide a full expression for 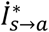 in the SI (SI Eqn. 108), and we validate our expression for *ρ*^2^ using simulations (SI Fig. S5). An ideal sensor with no internal noise corresponds to *γ* → ∞. Taking this limit in Eqn. <u>5</u> results in the expression for 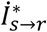 in Eqn. <u>3</u>. Conversely, internal noise degrades information about the signal, and the information rate becomes 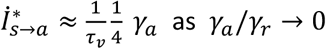. The derivation of 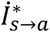 also provides the optimal kernel for constructing 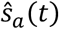, which we discuss in the SI (section “Optimal kernel for estimating signal from kinase activity”).

### Single-cell measurements constrain signal and kinase properties

To quantify the information rates above, we then performed single-cell tracking and FRET experiments to measure the parameters characterizing the signal statistics, kinase response function, and kinase noise statistics. As the attractant, we used aspartate (Asp), to which the *E. coli* chemotaxis signaling pathway responds with the highest sensitivity among known attractants (82).

To quantify the signal statistics, we recorded trajectories of cells swimming in multiple background concentrations of Asp: *c*_0_ = 0.1, 1, and 10 μM (Fig. 2A). Single cells in the clonal population exhibited a range of phenotypes (79,83–91). Therefore, as before (17), we focused on a typical cell by estimating the median single-cell parameter values in the population. In particular, we binned cells by the fraction of time spent running, *P*_*run*_, and computed *V(t)* among cells with the median *P*_*run*_. The parameters 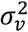 and τ_*v*_ in each background *c*_0_ were then estimated by fitting *V(t)* with a decaying exponential. These parameters depended weakly on *c*_0_, and their values in *c*_0_ = 1 μM were 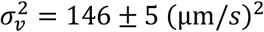 and τ_*v*_ = 1.1*9 ±* 0.01 *s* (see SI Fig. S1AB for all values).

**Figure 2.**
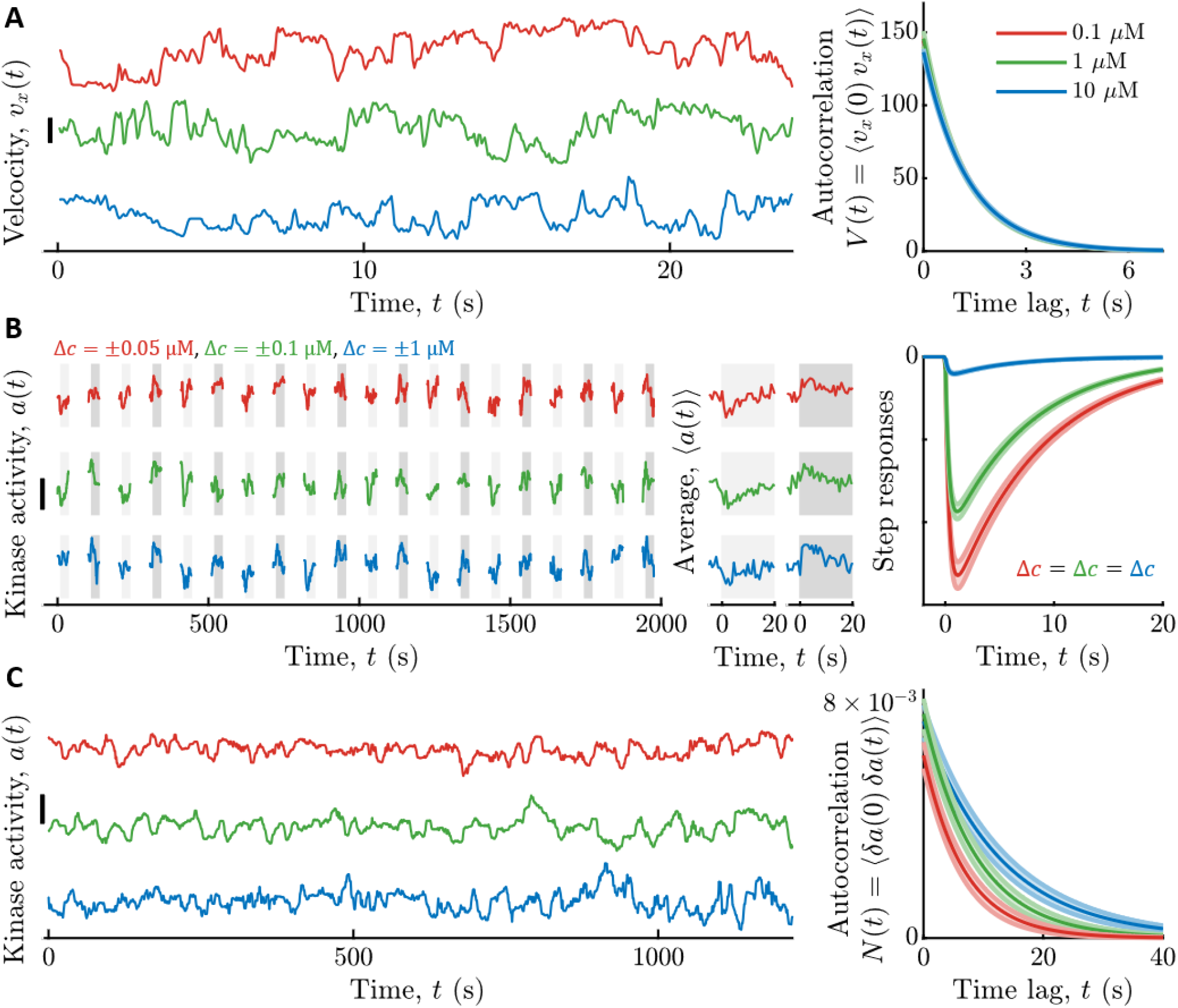
Measured signal statistics and kinase responses and fluctuations in different background ligand concentrations. **A)** Signal statistics. Left: Representative time series of up-gradient velocity *vx* from three individual cells are shown, one in each aspartate (Asp) concentration *c*0. Scale bar is 20 μm/s. Cells were binned by the fraction of time spent running, *Prun*, and the velocity autocorrelation function *V*(*t*) was computed by averaging over cells with the median *Prun*. The parameters of *V*(*t*) were extracted by fitting a decaying exponential to the data. Right: *V*(*t*) model fits for each *c*0. The curves are on top of each other. Vertical axis units are (μm/s)2. Throughout, shading is standard error of the mean (SEM), and line colors indicate *c*0: Red: 0.1 μM; Green: 1 μM; Blue: 10 μM. **B)** Linear responses. Left: Kinase activity was measured by FRET in blocks of 25 seconds, separated by 65 seconds without illumination. In each block, after 5 s, concentration was stepped up (light gray shading) or down (dark gray shading) around *c*0, then maintained for 20 s, then returned to *c*0. Concentration step sizes Δ*c* were different for each *c*0 (shown above the panel). Shown are three representative cells, one from each *c*0. Scale bar is 0.3. Middle: Average responses of the cells in the left panel to steps up (light gray) and steps down (dark gray). Single-cell responses were fit to extract parameters of the response function *Kr*(*t*). Right: Model fits for kinase responses to a steps size Δ*c*, using population-median parameters. The gain *Gr* decreases with *c*0. **C)** Noise statistics. Left: Fluctuations in kinase activity were measured in constant background concentrations. 9 Representative time series from three cells are shown, one from each *c*0. Scale bar height is 0.3. Parameters of the slow noise autocorrelation function were fit to single-cell traces using Bayesian filtering (17,102). Right: Estimated noise autocorrelation functions with population median parameters. Vertical axis units are kinase activity squared.

We measured kinase response functions as before (17), using a microfluidic device in which we can deliver controlled chemical stimuli with high time resolution (~100 ms) (50). Cells immobilized in the device were delivered ten small positive and negative step changes of Asp concentration around multiple backgrounds *c*_0_ (Fig. 2B). Kinase responses were measured in single cells through FRET (48–50,52,92–94) between CheZ-mYFP and CheY-mRFP1. Then we fit each cell’s average response to *K*_*r(t)*_above, and computed the population-median parameter values. Since τ_1_ estimated this way includes the relatively slow dynamics of CheY-CheZ interactions, we used τ_1_ = 0 for calculations below, which only slightly overestimates 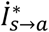. The adaptation time τ_2_ depended weakly on *c*_0_ (in *c*_0_ = 1 μM, τ_2_ = 7.4 *±* 0.3 *s*) (Fig. S1D), but *G*_*r*_ varied significantly: for *c*_0_ = {0.1, 1, 10} μM we measured 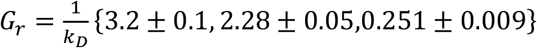 (Fig.S1EF).

The dependence of *G*_*r*_ on *c*_0_ was consistent with the Monod-Wyman-Changeux (MWC) model for kinase activity (29,95–97), which captures numerous experimental measurements (50,52,93,94,98). In particular, 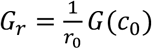 where 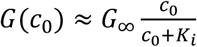 is the MWC gain, *K*_i_ is the dissociation constant of two-state receptors for Asp when in their inactive state, and *G*_∞_ is a constant (SI section “Modeling kinase activity”). Thus, in the “linear-sensing” regime (*c*_0_≪*K*_*i*_), the gain is constant,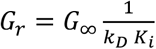, and in the “log-sensing” regime (*c*_0_ ≫ *K*_*i*_) (99–101), the gain decreases with background, 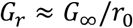. Fitting the measured *G*_*r*_ to the MWC model gave *G*_∞_ = 3.5 *±* 0.1 and *K*_*i*_ = 0.81 *±* 0.04 μM.

Finally, we estimated the parameters of slow kinase fluctuations by measuring kinase activity in single cells experiencing constant Asp concentrations *c*_0_ (Fig. 2C). The diffusivity *D*_*n*_ and time scale τ_*n*_ of these fluctuations were extracted from each time series using Bayesian filtering (17,102). We then computed the population-median parameter values. Both of these parameters depended weakly on *c*_0_, and their values in *c*_0_ = 1 μM were *D*_*n*_ = 8.1 *±* 0.*9* × 10^−4^ *s*^−1^ and τ_*n*_ = 8.7 *±* 0.*9 s* (see Fig. S1CD for all values).

### Comparing *E. coli* to the physical limit

Both *E. coli*’s information rate, 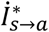, and the physical limit, 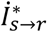, are proportional to *g*^2^ in shallow gradients. Therefore, using the measured parameters, we plotted the information rates per *g*^2^ as functions of *c*_0_ (Fig. 3A), for values of *g* in which we previously measured *E. coli*’s gradient-climbing speeds (17). Doing so reveals that *E. coli* are surprisingly far from the physical limit: in shallow gradients, 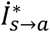 is at least two orders of magnitude below 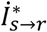 across all background concentrations.

**Figure 3.**
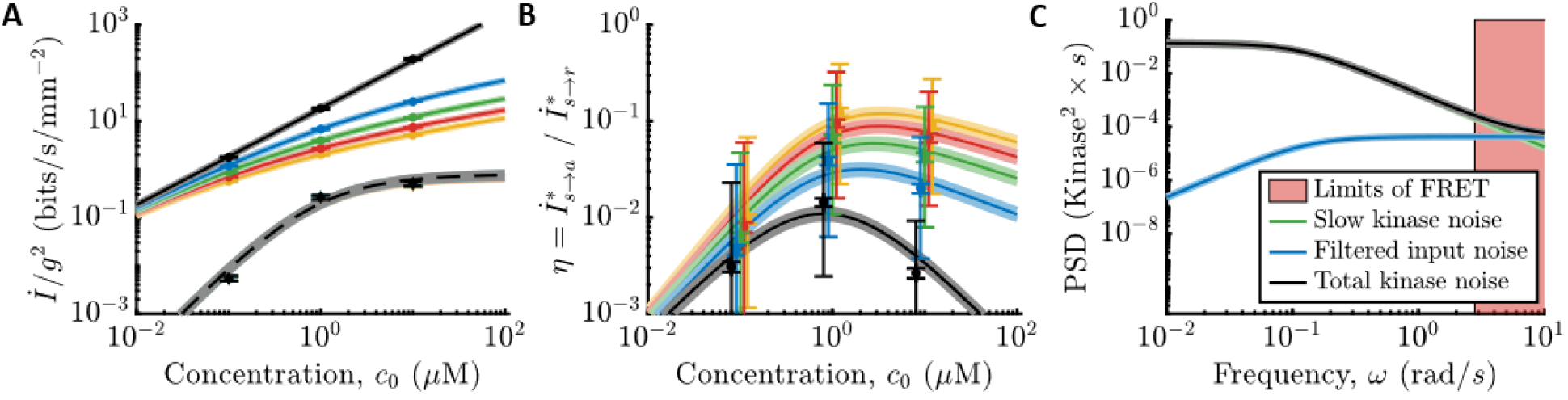
Comparing *E. coli*’s sensing accuracy to the physical limit. **A)** Information rates per gradient steepness squared, *g*2, in molecule arrival rate, 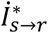 (SI Eqn. 46; solid lines), and in kinase activity, 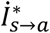 (SI Eqn. 108; dashed lines use the MWC model gain *G*(*c*0) and remaining parameters measured in *c*0=1 μM) for gradients of varying steepness, *g*∈{0+,0.1,0.2,0.3,0.4} mm–1 in black, blue, green, red, yellow, where 0+ is the limit of an infinitely shallow gradient. Dots are experimental measurements. Error bars and shading are the SEM. *E. coli* are far from the physical limit when signals are weak and sensor quality matters. **B)** 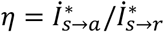 versus *c*0. Colors and markers are the same as in (A). Shading and small error bars on the dots are the SEM. Large error bars on the dots are estimates of 95% confidence intervals of population variation in η, assuming that swimming, kinase response, and kinase noise parameters are uncorrelated. Dots are shifted slightly for visual clarity. **C)** Fit models for the PSD’s of noise sources in *c*0=1 μM. Green: Slow noise in kinase activity. Blue: Molecule arrival noise filtered through the kinase response function. Black: Sum of green and blue. Red shading: Experimentally-inaccessible time scales using CheY-CheZ FRET. See also SI Fig. S3 and the SI section “Modeling kinase activity.”

To quantify this comparison, we computed the ratio of *E. coli*’s information rate and the physical limit,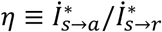 (Fig. 3B, small error bars). In vanishingly small gradients (black curve), η is independent of *g*. In this regime, 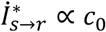 in all background concentrations, and the shape of η is determined by the gain of kinase response, *G*_*r*_. When *c*_0_ ≪ *K*_*i*_, the gain is constant, and η increases with background, η ∝ *c*_0_. When *c*_0_ ≫ *K*_*i*_, *G*_*r*_ decreases and cancels out increasing *c*_0_, so η ∝ 1*/c*_0_. These two regimes are separated by a peak at *c*_0_ = *K*_*i*_, where η ≈ 0.014 *±* 0.002 at our closest measurement. As the gradient gets steeper, η increases, up to η ≈ 0.1 when *g* = 0.4 mm^−1^. This larger value of η does not mean that *E. coli* count nearly every molecule in steeper gradients. Instead, the physical limit saturates (solid lines decreasing with *g* in Fig. 3A). Thus, in a steep gradient, even a poor sensor can infer the signal with decent accuracy.

Although typical cells in a population are far from the sensing limit, individual cells exhibit non-genetic diversity in sensing and swimming phenotypes (49,50,52,83,90,98), which could cause a significant fraction of the population to approach the limit. Our experimental setup did not allow us to measure all parameters in the same single cells, limiting our ability to answer this question. However, we do have single-cell parameters from different cells. Assuming that swimming, kinase responses, and kinase noise parameters are uncorrelated across cells, we use a maximum-likelihood approach to estimate the variability of η in the population (SI section “Estimating population variability in η”). This analysis indicates that although the 95th percentile of the population can be ~5 times closer to the physical limit, they are still far from it (Fig. 3B, large error bars).

In Fig. 3C, we show the power spectral density (PSD) of slow noise in kinase activity (green line) compared to the PSD of filtered molecule arrival noise (blue line) in *c*_0_ = 1 μM. If *E. coli* were close to the physical limit, nearly all noise in kinase activity would come from filtered molecule arrivals. Instead, slow kinase fluctuations are much larger over the range of frequencies observable in the experiment (Fig. 3C, outside the pink region). Thus, *E. coli*’s chemosensing is limited by constraints on its internal signal processing, rather than the external physics of ligand diffusion.

In Fig. 4, we demonstrate what this means for *E. coli* by simulating run-tumble motion in a gradient and constructing the optimal signal estimates (see SI section “Simulation details”). The top panels of Fig. 4A show the observed quantities: molecule arrival rate for an ideal cell, and kinase activity for *E. coli*. The bottom panels show the optimal estimates of the signal in each case, 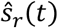 and 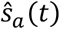, overlaid on the true signal. The estimate from kinase activity, 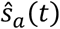, is visibly lower-quality than 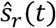. Quantitatively, 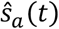 is less correlated with the true signal by nearly a factor of 10, and likewise kinase activity encodes about 10 times less information about signals than the physical limit. This figure shows the best-case scenario among those in Fig. 3AB; in shallower gradients or other background concentrations, this discrepancy increases to 100-fold or more.

**Figure 4.**
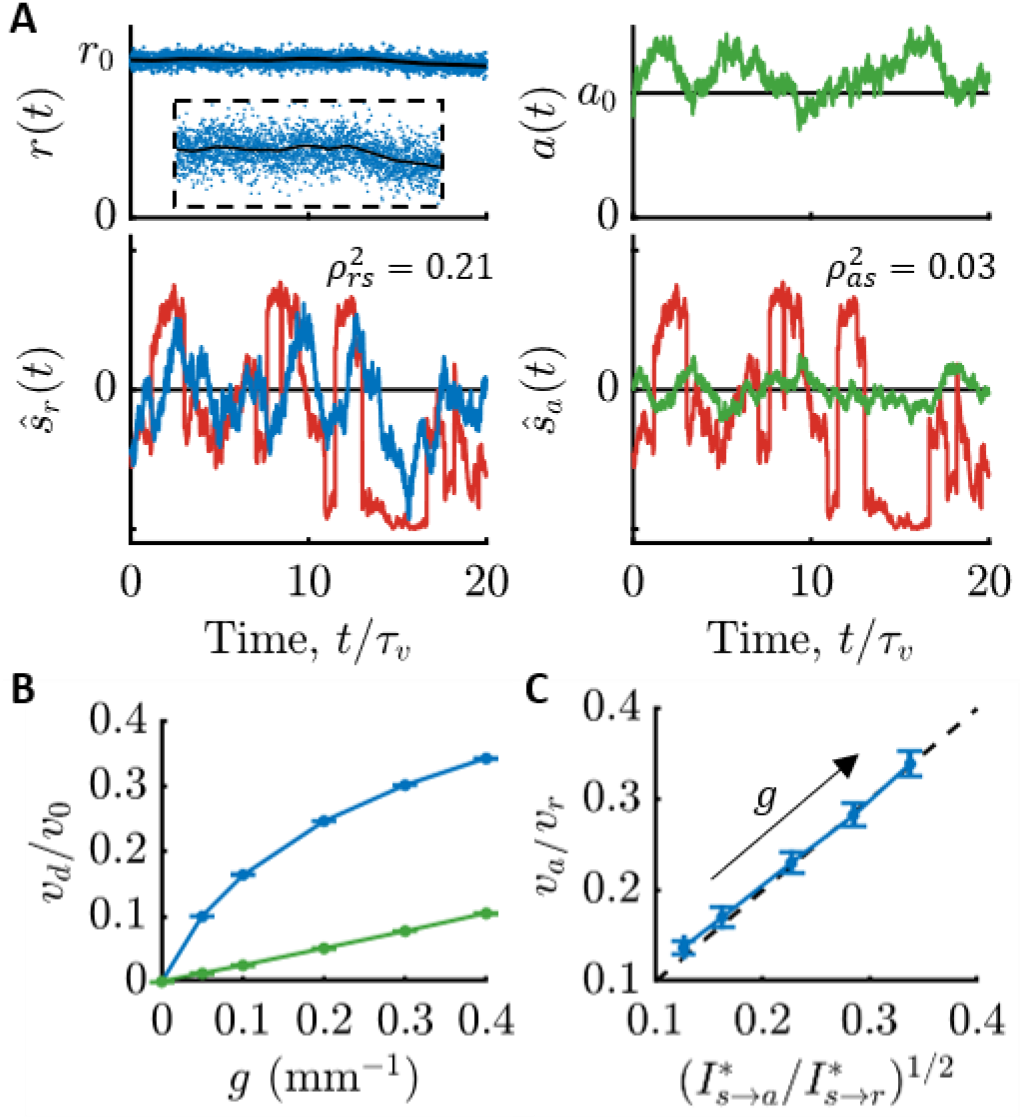
Consequences of *E. coli* being far from the physical limit on sensing. **A)** Simulation of run-tumble motion in a concentration gradient and optimal signal estimates using measured parameters (Fig. S1; *c*0=1 μM, *g*=0.4 mm^−1^). Top-left: An ideal cell directly observes molecule arrival rate *r*(*t*) (blue dots). Black line is the mean, ⟨*r*(*t*)⟩=k*D c*(*t*). Inset is the entire trajectory zoomed in to see the subtle changes in *c*(*t*). Bottom-left: Optimal signal estimate from molecule arrivals, 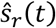 (blue), overlaid on the true signal, *s*(*t*) (red). Top-right: Simulated *E. coli* respond to molecule arrivals with changes in kinase activity (green). Bottom-right: Optimal signal estimate from kinase activity, 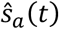 (green), overlaid on the true signal, *s*(*t*) (red). 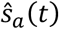 is visibly lower-quality than 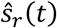. Squared Pearson correlation coefficients, 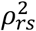 and 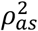, between the estimates and the true signal in each bottom panel quantify their accuracy. **B)** Chemotactic drift speed normalized by swimming speed, *v_d_*/*v*_0_, as a function of gradient steepness, *g*, for ideal cells (blue) and *E. coli* (green) in simulations (*c*0=1 μM; SI section “Simulation details”). Error bars in (B) and (C) are SEMs. **C)** Information lost between particle arrivals and kinase activity causes *E. coli* to climb gradients at speeds, *v_a_*, that are smaller than those of ideal cells, *v_r_*, by a factor of 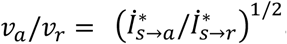. Thus, *E. coli* likely climb gradients slower than the physical limits of sensing allow. Dots are *g*={0.05,0.1,0.2,0.3,0.4} mm–1, from top-right to bottom-left.

Since information is needed for chemotaxis, this result implies that an ideal cell with the same swimming speed and run duration as *E. coli* (e.g. same *γ*_*r*_) could climb gradients much faster than *E. coli*. To support this, we simulated chemotaxis of ideal cells and *E. coli*-like cells in gradients of varying steepness. Fig. 4B indeed shows that ideal cells (blue), which directly observe particle arrival rate *r*, climb gradients much faster than *E. coli*-like cells (green), which only have access to kinase activity *a*. In Fig. 4C, we trace this reduction in drift speed directly back to *E. coli*’s loss of behaviorally-relevant information compared to an ideal cell. Our previous theory (17) predicts that the ratio of the *E. coli* cells’ drift speed, *v*_*a*_, to the ideal cell’s drift speed, 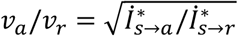, and plotting these ratios against each other in Fig. 4C shows good agreement. Thus, being far from the limits of chemosensing likely has a dramatic impact on *E. coli*’s gradient-climbing performance, especially in shallow gradients.

## Discussion

Living systems process information to perform survival-relevant functions, suggesting selection might optimize information processing. Here, we asked whether chemotaxing *E. coli* approach the physical limits on information processing set by counting diffusing ligand molecules. To make this question experimentally-testable, we devised a general approach that asks: how accurately can behaviorally-relevant signals be inferred from a cell’s interval variables (kinase activity) compared to the first physically-measureable quantity (ligand molecule arrivals). Accuracy in each case was quantified by an information rate, which we derived analytically. Then, we quantified these information rates in experiments using single-cell FRET measurements of cells’ CheA kinase activity in multiple background concentrations. Our results show that *E. coli* encode far less information than the physical limit, and thus *E. coli*’s chemosensing is shaped by internal constraints rather than the physical limit. The functional implication is that *E. coli* likely climb chemical gradients much slower than the physical limit on chemosensing allows. Thus, with the same signal-to-noise of particle arrivals, *γ*_*r*_ (set by the swimming speed, run duration, background concentration, and gradient steepness), in principle it may be possible to evolve or engineer a microswimmer that would climb gradients much faster than *E. coli*.

Our results are contrary to the belief, held in the field for nearly 50 years, that *E. coli*’s chemosensing approaches the physical limit, dramatically revising our understanding of bacterial chemotaxis. Since Berg and Purcell did not have direct access to *E. coli*’s uncertainty about ligand concentration, their argument for *E. coli*’s optimality assumed that cells must estimate the change in concentration over a *single* run, *Δc*, with uncertainty less than *Δc* (Eqn. 57 in Ref. (4)). Using experimental measurements and their physical limit, they computed the minimum required averaging time, *T*, for this condition to be met if the cell had access to particle arrivals. They found that measured bacterial run durations were slightly longer than the minimum *T*, and argued that chemotaxis would be impossible with shorter runs. Thus, they concluded that the bacterial chemotaxis machinery is nearly optimal. The problem with this argument is its first assumption: that in order to climb gradients, *E. coli*’s sensing machinery must exceed a stringent signal-to-noise threshold, so as to accurately infer the gradient direction in each run. Instead, *E. coli*’s displacement along the gradient accumulates their inferences over many runs. Therefore, even when individual tumble decisions are inaccurate, cells still climb the gradient on average, with no hard threshold on accuracy. In fact, we can show that Berg and Purcell’s assumption is too stringent: in our notation, their threshold condition can be written as *γ*_*r*_ > 16*/*3 (SI section “Berg & Purcell’s SNR threshold for chemotaxis”), but both the ideal cells and the *E. coli* cells simulated in Fig. 4B are able to climb the gradient when *g* = 0.05 mm^−1^, *c*_0_ = 1 μM, and *γ*_*r*_ = 0.15 ≪ 16*/*3.

Our results also disagree with those of Ref. (62), which argued that the marine bacterium *Vibrio ordalii* senses chemical signals with accuracy within a factor of ~6 of the physical limit, based on fits of agent-based simulations to measurements of bacteria climbing dynamic chemical gradients. We believe the reason for this difference is that their model assumed cells infer *s(t)* in independent time windows of duration *T* = 0.1 *s*. However, the signal is correlated over a time τ_*v*_ > *T*, so an ideal agent can average out molecule arrival noise for times up to τ_*v*_. This increases the theoretical limit, and thus *V. ordalii*’s distance from it, by a factor of 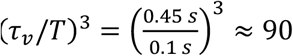, due to the 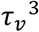 in *γ*_*r*_ (Eqn. <u>3</u>) (related to the *T*^3^ in Ref (35)). This suggests that chemosensing in other bacterial species, besides *E. coli*, may also be internally-limited. Beyond bacteria, our results call for a recalibration of expectations about the extent to which biology approaches physical limits on information processing or other functions.

Why are *E. coli* so far from the physical limit? One possibility is the physical implementation of their sensory system may impose trade-offs. For example, the need to operate over a wide range of background concentrations (99–101) suppresses response gain in high backgrounds, but the noise stays constant, reducing information. Cells may need to amplify signals above downstream noise sources, such as stochastic motor switching, requiring the densely-packed arrays seen universally across bacterial species (103), but strongly-coupled CheA kinases likely also introduce noise. Indeed, the dense localization of receptors suggests that molecule counting is not limiting, since if it were, the optimal strategy would be to uniformly distribute the receptors (4). *E. coli* also need to sense amino acids, sugars, and peptides (82,104) with different receptors, but the presence of multiple receptor types in the array reduces the response to any one ligand (94). Another possibility is that *E. coli* may be, and likely are, under selection pressures to perform other tasks, such as localize at concentration peaks (76,77,105,106). Laboratory strains have long been selected for chemotaxis via collective migration assays (107–109). The steep gradients generated during migration, reaching *g* ≈ 1 mm^−1^ or steeper (110–112), might obviate the need for a high-fidelity sensor. Lastly, increasing information about signals might be possible, but too costly in resources or energy to be worth the gain in fitness (20–26,32,113–115). The mechanism of amplification is not well understood, but recent work has argued that it consumes energy (116–118). These possibilities might be distinguished by measuring information rates of single cells in an isogenic population or information rates of mutants. If any single cell approaches the physical limit, it would mean that *E. coli* are not limited by hard implementation constraints, but rather by costs or competing objectives. Answering this question will likely inform our thinking about the relevant physical constraints on information processing in other systems.

Physical limits, and whether biology approaches them, have long inspired physicists’ curiosity (1–17). At first glance, our results seem to call into question the value of normative theories and physical limits for understanding biological information processing. However, our findings were only possible because we derived a physical limit that provided a reference point against which to compare. At the same time, our results motivate going beyond physical limits and taking seriously the system-specific, physical and biological *constraints* on biological information processing. Going forward, we expect *E. coli* chemotaxis will be a valuable template for studying physical limits and constraints on information processing in higher organisms.

## Methods

### Strains and plasmids

All strains and plasmids used are the same as in our recent work (17). The strain used for the FRET experiments is a derivative of *E. coli* K-12 strain RP437 (HCB33), a gift of T. Shimizu, and described in detail elsewhere (49,50). The FRET acceptor-donor pair (CheY-mRFP and CheZ-mYFP) is expressed in tandem from plasmid pSJAB106 (49) under an isopropyl β-D-thiogalactopyranoside (IPTG)-inducible promoter. The glass-adhesive mutant of FliC (FliC*) was expressed from a sodium salicylate (NaSal)-inducible pZR1 plasmid (49). The plasmids are transformed in VS115, a *cheY cheZ fliC* mutant of RP437 (49) (gift of V. Sourjik). RP437, the direct parent of the FRET strain and also a gift from T. Shimizu, was used to measure swimming statistics parameters. All strains are available from the authors upon request.

### Cell preparation

Single-cell FRET microscopy and cell culture was carried out essentially as described previously (17,49,50,52). Cells were picked from a frozen stock at −80°C and inoculated in 2 mL of Tryptone Broth (TB; 1% bacto tryptone, 0.5 % NaCl) and grown overnight to saturation at 30°C and shaken at 250 RPM. Cells from a saturated overnight culture were diluted 100X in 10 mL TB and grown to OD600 0.45-0.47 in the presence of 100 μg/ml ampicillin, 34 μg/ml chloramphenicol, 50 μM IPTG and 3 μM NaSal, at 33.5°C and 250 RPM shaking. Cells were collected by centrifugation (5 min at 5000 rpm, or 4080 RCF) and washed twice with motility buffer (10 mM KPO4, 0.1 mM EDTA, 1 μM methionine, 10 mM lactic acid, pH 7), and then were resuspended in 2 mL motility buffer, plus the final concentration of Asp. Cells were left at 22°C for 90 minutes before loading into the microfluidic device. All experiments, FRET and swimming, were performed at 22-23°C.

For swimming experiments, cells were prepared similarly. Saturated overnight cultures were diluted 100X in 5 mL of TB. After growing to OD600 0.45-0.47, 1 mL of cell suspension was washed twice in motility buffer with 0.05% w/v of polyvinylpyrrolidone (MW 40 kDa) (PVP-40). Washes were done by centrifuging the suspension in an Eppendorf tube at 1700 RCF (4000 RPM in this centrifuge) for 3 minutes. After the last wash, cells were resuspended with varying background concentrations of Asp.

### Microfluidic device fabrication and loading for FRET measurements

Microfluidic devices for the FRET experiments (50,52,92) were constructed from polydimethylsiloxane (PDMS) on 24 x 60 mm cover glasses (#1.5) following standard soft lithography protocols (119), exactly as done before (17).

Sample preparation in the microfluidic device was conducted as follows. Five inlets of the device were connected to reservoirs (Liquid chromatography columns, C3669; Sigma Aldrich) filled with motility buffer containing various concentrations of Asp through polyethylene tubing (Polythene Tubing, 0.58 mm id, 0.96 mm od; BD Intermedic) (see SI of (17)). The tubing was connected to the PMDS device through stainless steel pins that were directly plugged into the inlets or outlet of the device (New England Tubing). Cells washed and suspended in motility buffer were loaded into the device from the outlet and allowed to attached to the cover glass surface via their sticky flagella by reducing the flow speed inside the chamber. The pressure applied to the inlet solution reservoirs was controlled by computer-controlled solenoid valves (MH1; Festo), which rapidly switched between atmospheric pressure and higher pressure (1.0 kPa) using a source of pressurized air. Only one experiment was conducted per device. *E. coli* consume Asp, so all experiments below were performed with a low dilution of cells to minimize this effect. The continuous flow of fresh media also helped ensured that consumption of Asp minimally affected the signal cells experienced.

### Single-cell FRET imaging system

FRET imaging in the microfluidic device was performed using the setup as before (17), on an inverted microscope (Eclipse Ti-E; Nikon) equipped with an oil-immersion objective lens (CFI Apo TIRF 60X Oil; Nikon). YFP was illuminated by an LED illumination system (SOLA SE, Lumencor) through an excitation bandpass filter (FF01-500/24-25; Semrock) and a dichroic mirror (FF520-Di02; Semrock). The fluorescence emission was led into an emission image splitter (OptoSplit II; Cairn) and further split into donor and acceptor channels by a second dichroic mirror (FF580-FDi01-25×36; Semrock). The emission was then collected through emission bandpass filters (F01-542/27-25F and FF02-641/75; Semrock) by a sCMOS camera (ORCA-Flash4.0 V2; Hamamatsu). RFP was illuminated in the same way as YFP except that an excitation bandpass filter (FF01-575/05-25; Semrock) and a dichroic mirror (FF593-Di03; Semorock) were used. An additional excitation filter (59026x; Chroma) was used in front of the excitation filters. To synchronize image acquisition and the delivery of stimulus solutions, a custom-made MATLAB program controlled both the imaging system (through the API provided by Micro-Manager (120)) and the states of the solenoid valves.

### Computing FRET signal and kinase activity

FRET signals were extracted from raw images using the E-FRET method (121), which corrects for different rates of photobleaching between donor and acceptor molecules. In this method, YFP (the donor) is illuminated and YFP emission images (*I*_*DD*_) and RFP (the acceptor) emission images (*I*_*DA*_) are captured. Periodically, RFP is illuminated and RFP emission images are captured (*I*_*AA*_). From these, photobleach-corrected FRET signal is computed as before (17), which is related to kinase activity *a(t)* by an affine transform when CheY and CheZ are overexpressed (17,93). All parameters associated with the imaging system were measured previously (17).

In each experiment, we first delivered a short saturating stimulus (1 mM MeAsp plus 100 µM serine (94)) to determine the FRET signal at minimum kinase activity, followed by motility buffer with Asp at background concentration *c*_0_. Before the saturating stimulus was delivered, the donor was excited every 0.5 seconds to measure *I*_*DD*_ and *I*_*DA*_ (see SI of (17)) for 5 seconds. Then the stimulus was delivered for 10 seconds, and the donor was excited every 0.5 seconds during this time. Before and after the donor excitations, the acceptor was excited three times in 0.5-second intervals to measure *I*_*AA*_ (see SI of (17)). After the stimulus was removed, the acceptor was excited three more times at 0.5-second intervals. Imaging was then stopped and cells were allowed to adapt to the background for 120 seconds.

Stimulus protocols for measuring kinase linear response functions and fluctuations are described below. At the end of each experiment, we delivered a long saturating stimulus (1 mM MeAsp plus 100 µM serine) for 180 seconds to allow the cells to adapt. Then we removed the stimulus back to the background concentration, eliciting a strong response from the cells, from which we determined the FRET signal at maximum kinase activity. The donor was excited for 5 seconds before the saturating stimulus and 10 seconds after it, every 0.5 seconds. Before and after these donor excitations, the acceptor was excited three times in 0.5-second intervals. The cells were exposed to the saturating stimulus for 180 seconds. The donor was excited every 0.5 seconds for 5 seconds before cells were exposed to motility buffer with Asp at background concentration *c*_0_, followed by 10 seconds of additional donor excitations. Before and after the donor excitations, the acceptor was again excited three times in 0.5-second intervals.

FRET signals were extracted as before (17). The FRET signal at minimum kinase activity, *FRET*_*min*_, was computed from the average FRET signal during the first saturating stimulus. The FRET signal at maximum kinase activity, *FRET*_*max*_, was computed from the average FRET signal during the first quarter (2.5 seconds) of the removal stimulus at the end of the experiment. Kinase activity was then computed from corrected FRET signal: 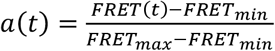.

### Kinase linear response functions

Experiments were performed in Asp background concentrations *c*_0_ of 0.1, 1, and 10 µM. Measurements were made in single cells, and at least three replicates were performed per background. FRET level at minimum kinase activity was measured at the beginning of each experiment, as described above. After this, a series of stimuli were delivered to the cells in the microfluidic device. Cells were only illuminated and imaged when stimulated in order to limit photobleaching. Before each stimulus, cells were imaged for 7.5 seconds in the background concentration *c*_0_. Then, the concentration of Asp was shifted up to *c*_+_ > *c*_0_ for 30 seconds and imaging continued. Donor excitation interval was 0.75 seconds and acceptor excitations were done before and after the set of donor excitations. After this, imaging was stopped and the Asp concentration returned to *c*_0_ for 65 seconds to allow cells to adapt. Then, the same process was repeated, but this time shifting Asp concentration down to *c*_−_ < *c*_0_. Alternating up and down stimuli were repeated 10 times each. *c*_+_ and *c*_−_ varied with each experiment and each background *c*_0_. Finally, FRET level at maximum kinase activity was measured at the end of each experiment, as described above. The whole imaging protocol lasted <2200 seconds. In total, cells spent <60 minutes in the device, from loading to the end of imaging.

These data were analyzed as before (17) to extract linear response parameters for each cell. In brief, the responses of a cell to all steps up or steps down in concentration were averaged and the standard error of the response at each time point computed. Model parameters were extracted by maximizing the posterior probability of parameters given data, assuming a Gaussian likelihood function and log-uniform priors for the parameters. The uncertainties of single-cell parameter estimates were generated by MCMC sampling the posterior distribution. Finally, the population-median parameters were computed from all cells in experiments in a given background *c*_0_. Uncertainty 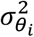 of the population-median value of parameter *θ*_*i*_, with *θ* = *(G*, τ_1_, τ_2_), was computed using:

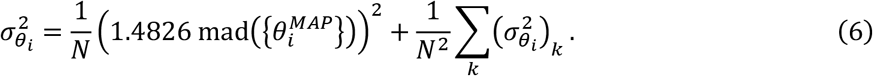

This expression accounts both for cell-to-cell variations (first term) and uncertainties in the single-cell estimates (second term). *N* is the number of cells. 1.4826 mad() is an outlier-robust uncertainty estimate that coincides with the standard deviation when the samples are Gaussian-distributed, and mad() is the median absolute deviation, used previously (17). 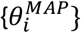 are the single-cell maximum *a-posteriori* (MAP) estimates of parameter 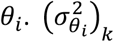 is the uncertainty of 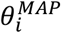 in cell k, which was computed using

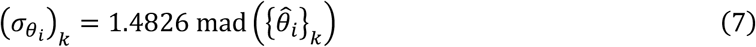

where 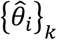 are the samples from the kth cell’s posterior via Markov Chain Monte Carlo (MCMC).

### Fitting the MWC kinase gain

Parameters *G*_∞_ and *K*_*i*_ of the MWC model gain were estimated by fitting the model to estimated values of *G* in each background *c*_0_. The fit was done by minimizing the sum of squared errors between the logarithms of the measured *G* and predicted values of *G*. Since the estimated values of *G* varied by about an order of magnitude, taking the logarithms ensured that the smallest value of *G* had similar weight as largest value in the objective function.

### Statistics of noise in kinase activity

Fluctuations in kinase activity were measured in the same Asp background concentrations *c*_0_ as above, as well as *c*_0_ = 0 μM. At least three replicate experiments were performed per background. FRET level at minimum kinase activity was measured at the beginning of each experiment, as described above. After these measurements, imaging was then stopped and cells were allowed to adapt to the background for 120 seconds. After this, cells were imaged for about 1200 seconds. Throughout, donor excitations were done every 1.0 second, except when it was interrupted by acceptor excitations, which were conducted every 100 donor excitations (see SI of (17)). Finally the FRET level at maximum kinase activity was measured at the end of each experiment, as described above. The whole imaging protocol lasted <1400 seconds. In total, cells spent about < 60 minutes in the device, from loading to the end of imaging.

These data were analyzed as before (17). Bayesian filtering methods (102) were used to compute the likelihood of the parameters given the data, and the prior distribution was taken to be uniform in log. Single-cell estimates and uncertainties of the noise parameters were extracted from the posterior distribution as described above. In each background *c*_0_, the population median parameter values were computed, and their uncertainties were computed as described above, with *θ* = *(D*_*n*_, τ_*n*_).

### Swimming velocity statistics

Cells were prepared and imaged as before (17). After the second wash step of the <u>Cell preparation</u> section above, cells were centrifuged again and resuspended in motility buffer containing a background concentration of Asp *c*_0_. The values of *c*_0_ used here were the same as in the FRET experiments, including *c*_0_ = 0 μM. Then, the cell suspension was diluted to an OD600 of 0.00025. This low dilution of cells both enables tracking and minimizes the effect of cells consuming Asp. The cell suspension was then loaded into µ-Slide Chemotaxis devices (ibidi; Martinsried, Germany). Swimming cells were tracked in one of the large reservoirs. 1000-s movies of swimming cells were recorded on a Nikon Ti-E Inverted Microscope using a CFI Plan Fluor 4X objective (NA 0.13). Images were captured using a sCMOS camera (ORCA-Flash4.0 V2; Hamamatsu). Four biological replicates were performed for each background *c*_0_.

Cell detection and tracking were carried out using the same custom MATLAB as we used previously (17), with the same analysis parameters (see SI of that paper for details). Tumble detection was also carried out identically as before (17). There was no minimum trajectory duration, but cells were kept only if at least two tumbles were detected in their trajectory. For each cell, we computed the fraction of time spent in the “run” state *P*_*run*_. Then we constructed the distribution of *P*_*run*_, correcting for biases caused by the different diffusivities of cells with different *P*_*run*_ (17). As before (17), we then computed the correlation function of velocity along one spatial dimension for each cell, *V*_*i*_*(t)* = ⟨*v*_*x*_*(t*′*)v*_*x*_*(t*′ + *t)*⟩_*t*_′ among cells with *P*_*run*_ within *±*0.01 of the population-median value,. Finally, we computed a weighted average of the correlation functions over all cells in the population-median bin of *P*_*run*_, where trajectories were weighted by their duration, giving *V(t)*. In each background *c*_0_, for the median bin of *P*_*run*_, the average trajectory duration was ~7.6 seconds, and the total trajectory time was ≥ 2.7 × 10^4^ seconds.

These correlation functions *V(t)* in each background *c*_0_ and each experiment were fit to decaying exponentials 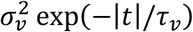 exp(−|*t*|*/*τ_*v*_ ), and the parameters and their uncertainties were extracted in two steps. First, we determined the MAP estimates of the parameters. An initial estimate of the parameters were esimated using the MATLAB *fit* function to fit exponentials to the *V(t)* in the time rang *t* ∈ [2 *Δt*, 10 s], with *Δt* = 50 ms. The estimated τ_*v*_ was used to get the uncertainty of *V(t)* in each experiment, as done before (17). Assuming a Gaussian likelihood function and parameters distributed uniformly in logarithm, the posterior distribution of parameter was constructed. In each experiment, MAP estimates of the parameters were extracted as done for the kinase parameters, and parameter uncertainties were computed from MCMC samples of the posterior distribution as above. Finally, we computed the average parameters 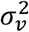 and τ_*v*_ over experimental replicates, as well as their standard errors over replicates.

### Additional error analysis

Once the variance of the population-median value of parameter *i* was computed, 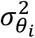, we propagated the uncertainty to functions of those parameters. For some function of the parameters, *f(θ)*, we computed the variance of *f(θ)*, 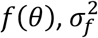, as:

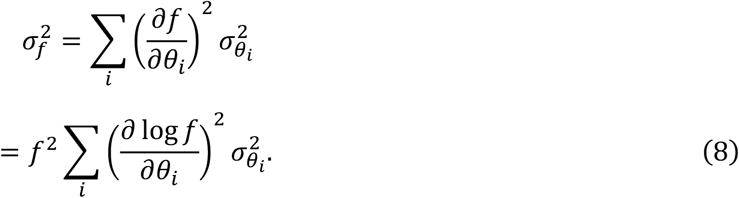

The equations above neglect correlations in the uncertainties between pairs of parameters. This was used to compute the uncertainties of 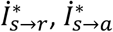, and η. The same formula was used to compute uncertainties of functions of time by applying the formula above pointwise at each time delay *t* and neglecting correlations in uncertainties between time points.

## Funding

This work was supported by the Alfred P. Sloan Foundation under grant G-2023-19668 (HM, TE, BB); by NIH awards R01GM106189 (TE), R01GM138533 (TE), R35GM158058 (TE), and R35GM138341 (BM); by Simons Investigator Award 624156 (BM); by the JST PRESTO grant JPMJPR21E4 (KK); and by the NSTC grant 112-2112-M-001-080-MY3 (KK). HM was supported by the Simons Foundation. KK was also supported by the Institute of Molecular Biology, Academia Sinica.

## Author contributions

BM and HM conceived the project. KK, HM, TE, and BM designed the experiments. KK, JO, RK, and HM performed the experiments. HM and KK analyzed the data. HM and BM derived the theoretical results. HM wrote the first draft of the manuscript. HM, BM, KK, and TE edited the manuscript.

## Competing interests

The authors declare no competing interests.

## Data availability

Source data for the main text figures will be provided online with the manuscript. Source data for the Supplementary Figures are contained in a Supplementary Data file.

## Code availability

Code to reproduce the main text figures will be available with the source data. All algorithms used are described in detail in the Supplementary Information.

## Notes

### Competing Interest Statement

The authors have declared no competing interest.

### Summary of Updates

Major edits to the text, new figures 1 and 4.

