## Supplemental information for "Chemotaxing *E. coli* do not count single molecules"

### Supplementary Text

#### Drift speed and information rate

We recently demonstrated that a cell's drift speed $v_d$ is set by the transfer entropy rate, $\dot{I}^*_{s\to m}$, from current signal $s(t) = \frac{1}{c_0}\frac{dc}{dt}$ to (the trajectory of) swimming behavior $m(t)$ (1). The transfer entropy rate from current signal to swimming behavior is defined as:

$$\dot{I}^*_{s\to m} = \lim_{dt\to 0} \frac{1}{dt} I(m(t+dt); s(t)|\{m(t)\}) \tag{1}$$

$$= \lim_{dt\to 0} \frac{1}{dt} \left\langle \log\left(\frac{P(m(t+dt)|s(t), \{m(t)\})}{P(m(t+dt)|\{m(t)\})}\right)\right\rangle. \tag{2}$$

Here, curly brackets denote the entire past of a variable, up to and including time $t$. Angled brackets indicate an average over the joint distribution of $s(t)$, past $m(t)$, and $m(t+dt)$. This quantifies how strongly the swimming transition probabilities depend on the current signal.

The transfer entropy rate from current signal determines the cell's drift speed (1):

$$\frac{v_d}{v_0} = \frac{(1-\alpha)\,\lambda_{R0}}{(1-\alpha)\,\lambda_{R0} + 2\,D_r} \left(\frac{2}{3}\frac{\dot{I}^*_{s\to m}}{\lambda_{R0}} P_{run}\right)^{1/2} \tag{3}$$

where $v_0$ is the cell's swimming speed, $\lambda_{R0}$ is the cell's average tumble rate, $\alpha$ is the persistence of the cell's orientation upon tumbling, $D_r$ is the rotational diffusion coefficient, and $P_{run}$ is the fraction of time the cell spends in the run state. This result comes from our previous demonstration that, in shallow gradients, both the drift speed and $\dot{I}^*_{s\to m}$ are proportional to the correlation between the cell's tumble rate $\lambda(t)$ and the signal $s(t)$. Since this correlation determines the cell's chemotaxis performance, $v_d/v_0$, $s(t)$ is the behaviorally-relevant signal.

#### Equivalence of transfer entropy and predictive information rates

Here we demonstrate that the transfer entropy rates above are equivalent to a predictive information rate, under some assumptions that are satisfied by bacterial chemotaxis. This relationship is useful because it allows us to derive expressions for the behaviorally-relevant information rates above.

Below, we will write transfer entropy rate from a signal $s(t)$ to a stochastic process $x(t)$, such as $r(t)$, $a(t)$, or $m(t)$. Starting from the definition above:

$$\dot{I}^*_{s\to x} = \lim_{dt\to 0} \frac{1}{dt} I(x(t+dt); s(t)|\{x(t)\}), \tag{4}$$

conditional mutual information can be written as a difference between two unconditioned mutual information terms:

$$= \lim_{dt\to 0} \frac{1}{dt} \Big( I\big(\{x(t+dt)\}; s(t)\big) - I\big(\{x(t)\}; s(t)\big)\Big). \tag{5}$$

This can be written as

$$= \left[\partial_T I\big(\{x(T)\}; s(t)\big)\right]_{T=t}. \tag{6}$$

Changing variables from $T$ to $\tau$, where $T = t + \tau$, we get:

$$= \left[\partial_\tau I\big(\{x(t+\tau)\}; s(t)\big)\right]_{\tau=0}. \tag{7}$$

Next, we use time stationarity to shift time $t$ by $-\tau$:

$$= \left[\partial_\tau I\big(\{x(t)\}; s(t-\tau)\big)\right]_{\tau=0}. \tag{8}$$

Finally, we can change variables to $\tau \to -\tau$, giving:

$$= -\left[\partial_\tau I\big(\{x(t)\}; s(t+\tau)\big)\right]_{\tau=0}. \tag{9}$$

This last step would not be allowed if the mutual information inside the time derivative was the entire past of $s$, i.e. $\{s(t+\tau)\}$.

Inside the time derivative above is the "predictive information" (2–4) between the entire past of the stochastic process $x(t)$ up to time $t$ and the signal $s(t)$ at some time $\tau$ into the future (if $\tau > 0$). The time derivative of this mutual information or predictive information is a monotonically decreasing function of $\tau$: the value of the signal $s$ at a time further in the future (larger $\tau$) becomes less correlated with past observations and thus harder to predict.

#### Data processing inequality

The causal structure of bacterial chemotaxis has a feedback loop: $m \to s \to r \to a \to m$. Tumbles affect the dynamics of $s$ by changing the cell's heading, which affect the dynamics of $r$, which affect $a$, which then feeds back to $m$ by modulating the tumble rate. In shallow gradients, this feedback is negligible, and transfer entropies obey a data processing inequality (5,6). Thus, information about the signal available in particle arrivals sets a fundamental upper limit on how much information a cell can get, which in turn sets an upper limit its gradient climbing speed:

$$\dot{I}^*_{s\to r} \geq \dot{I}^*_{s\to a} \geq \dot{I}^*_{s\to m} \propto (v_d/v_0)^2. \tag{10}$$

We further speculate that $\dot{I}^*_{s\to x}$ defined above (involving only the current $s(t)$) obeys a data processing inequality in bacterial chemotaxis, even when the gradient is not shallow. If the link from signal to motor behavior is severed, either by having kinases that don't respond to signals or motors that don't respond to kinase activity, then we argue that $\dot{I}^*_{s\to m} = 0$. In this case, as long as post-tumble headings have no preferred direction relative to the gradient direction, then knowing when the cell tumbled (contained in $\{m\}$) provides no information about the value of the current or future signal, $s(t+\tau)$. Therefore, in this case, $I\big(\{m(t)\}; s(t+\tau)\big) = 0$, and $\dot{I}^*_{s\to m} = 0$.

Then, the only way that the past of $m$ provides information about $s(t)$ is through chemotaxis responses to signals, which induce changes in run duration—longer than average runs imply $s(t) > 0$. But tumble behavior in response to signals passes through kinase activity $a$, so $a$ must be more informative than $m$ about $s(t)$.

#### Modeling concentration and molecule arrival rate

To derive the physical limit on behaviorally-relevant information for chemotaxis, we first need models for the dynamics of concentration $c(t)$, the signal $s(t) = \frac{d}{dt}\log\big(c(t)\big)$, and particle arrival rate $r(t)$. We consider a single cell navigating a shallow, static chemical gradient, $c(x) = c_0\, e^{g\,x} \sim c_0(1 + g\,x)$, that varies along one spatial dimension, $x$, in 3D space. In a static gradient, the signal is determined by the cell's motion in the gradient: $s(t) = \frac{d}{dt}\log(c) \approx \frac{1}{c_0}\frac{dc}{dt} = g\, v_x(t)$. As done before (1), we model the cell's up-gradient velocity, and thus the signal, as a Gaussian process with correlation function:

$$\langle s(t)\, s(t')\rangle = g^2\, V(t) = g^2\, \sigma_v^2 \exp\left(-\frac{|t - t'|}{\tau_v}\right). \tag{11}$$

Here, $\sigma_v^2 \approx \frac{v_0^2}{3} P_{run}$ is the variance of the cell's up-gradient velocity, $v_0$ is its swimming speed, and $P_{run}$ is the fraction of time it spends in the run state; and $\tau_v$ is the correlation time of the cell's velocity and the signal, $\tau_v^{-1} = (1 - \alpha)\, \lambda_{R0} + 2\, D_r$ , where $\lambda_{R0}$ is the cell's baseline tumble rate, $\alpha$ is the directional persistence, and $D_r$ is the rotational diffusion coefficient (1).

We model concentration and particle arrival rate as follows:

$$\frac{dc}{dt} = c_0\, s(t) \tag{12}$$

$$r(t) = k_D\, c(t) + \sqrt{r_0}\, \xi(t). \tag{13}$$

$k_D = 4\, D\, l$ is the diffusion-limited rate constant of particle arrivals to a membrane patch of radius $l$ and for ligand diffusion coefficient $D$ (7–9). The particle arrival noise obeys $\langle \xi(t)\, \xi(t')\rangle = \delta(t - t')$, and $r_0 = k_D\, c_0$ is the particle arrival rate in background concentration $c_0$. Although particle arrival events follow a Poisson process, this Gaussian approximation for the arrival rate is valid when the cell encounters many particles per run, $r_0\, \tau_v \gg 1$.

We will need several power spectral densities. To compute them, we take the Fourier transforms of Eqns. 12 and 13, and then solve for the Fourier transforms of $c(t)$ and $r(t)$, denoted $c(\omega)$ and $r(\omega)$:

$$c(\omega) = \frac{c_0}{\frac{\epsilon}{\tau_v} - i\,\omega} s(\omega) \tag{14}$$

$$r(\omega) = k_D\, c(\omega) + \sqrt{r_0}\, \xi(\omega). \tag{15}$$

Here we have introduced a small, dimensionless parameter $\epsilon \ll 1$ that we will take to zero later. Physically, this is as if the cell experiences a weak restoring force back to regions where concentration $c(x) = c_0$. Without it, the correlation function of $c(t)$, which is proportional to the cell's mean squared displacement, would diverge at long times. Everything else remains bounded and well-defined as $\epsilon$ goes to zero.

From these and the correlation function of $s(t)$, we derive the following power spectral densities:

$$S_s(\omega) = F[C_s(T)] = \frac{2\, g^2 \frac{\sigma_v^2}{\tau_v}}{\frac{1}{\tau_v^2} + \omega^2} \tag{16}$$

$$S_r(\omega) = F[C_r(T)] = \frac{r_0^2}{\frac{\epsilon^2}{\tau_v^2} + \omega^2} S_s(\omega) + r_0 \tag{17}$$

$$S_{rs}(\omega) = S_{sr}^*(\omega) = F[C_{rs}(T)] = \frac{r_0}{\frac{\epsilon}{\tau_v} + i\,\omega} S_s(\omega) \tag{18}$$

where $C_s(T) = \langle s(t)\, s(t+T)\rangle$ , $C_r(T) = \langle (r(t) - r_0)\,(r(t+T) - r_0)\rangle$ , and $C_{rs}(T) = \langle (r(t) - r_0)\, s(t+T)\rangle$, the Fourier transform is defined as $F[f(t)] = \int_{-\infty}^{\infty} f(t)\, e^{i\,\omega\, t}\, dt$, and the inverse transform defined as $F^{-1}[f(\omega)] = \frac{1}{2\pi}\int_{-\infty}^{\infty} f(\omega)\, e^{-i\,\omega\, t}\, d\omega$.

#### Total information in particle arrival rate is a trivial upper limit on information processing

To emphasize the importance of behaviorally-relevant information, we first derive the total information about past signals, $\{s(t)\}$, encoded in past particle arrival rate, $\{r(t)\}$:

$$\dot{I}_{s\to r} = \lim_{dt\to 0} \frac{1}{dt} I(r(t+dt); \{s(t)\}|\{r(t)\}) = \dot{I}(\{s\};\{r\}). \tag{19}$$

The last quantity is the mutual information rate (5,10,11) between $s$ and $r$, and the equality is valid when there is no feedback from $r$ to $s$ or in the regime of shallow gradients. We (1) and others (11) considered information quantities of this kind to quantify the total information about past signals encoded by *E. coli*'s chemotaxis signaling pathway (see also the section below, **Information about current versus past signals encoded in kinase activity**). The total information rate $\dot{I}_{s\to r}$ sets an upper limit on the information that *E. coli* can encode about signals. Computing it using the Gaussian approximations for $s$ and $r$, we find:

$$\begin{aligned}
\dot{I}_{s\to r} &= \lim_{\epsilon\to 0} \frac{-1}{4\pi}\int_{-\infty}^{\infty} \log\left(1 - \frac{|S_{rs}(\omega)|^2}{S_r(\omega)\, S_s(\omega)}\right) d\omega \\
&= \lim_{\epsilon\to 0} \frac{-1}{4\pi}\int_{-\infty}^{\infty} \log\left(1 - \frac{\frac{r_0^2}{(\epsilon/\tau_v)^2 + \omega^2} S_s(\omega)}{\frac{r_0^2}{(\epsilon/\tau_v)^2 + \omega^2} S_s(\omega) + r_0}\right) d\omega \\
&= \lim_{\epsilon\to 0} \frac{1}{4\pi}\int_{-\infty}^{\infty} \log\left(1 + \frac{r_0}{(\epsilon/\tau_v)^2 + \omega^2} S_s(\omega)\right) d\omega \to \infty.
\end{aligned} \tag{20}$$

Thus, we find that this total information rate provides a meaningless bound on *E. coli*'s information processing because it is dominated by information about past signals that are not relevant to chemotaxis.

#### Derivation of the physical limit on behaviorally-relevant information for chemotaxis

In this section we derive the information rate from current signal $s(t)$ to past particle counts $r$, which sets a fundamental upper limit on the information rate achievable by a cell. This information rate is given by the following transfer entropy rate:

$$\dot{I}^*_{s\to r} = \lim_{dt\to 0}\frac{1}{dt} I(r(t+dt); s(t)|\{r(t)\})$$
$$= -\left[\partial_\tau I\left(\{r(t)\}; s(t+\tau)\right)\right]_{\tau=0}. \quad (21)$$

The key quantity we need to derive is the mutual information inside of the derivative:

$$I\left(\{r(t)\}; s(t+\tau)\right) = \left\langle \log\left(\frac{P(s(t+\tau)|\{r(t)\})}{P\left(s(t+\tau)\right)}\right)\right\rangle = \left\langle \log\left(\frac{P\left(\{r(t)\} \mid s(t+\tau)\right)}{P(\{r(t)\})}\right)\right\rangle. \quad (22)$$

In general, it is difficult to derive the conditional distributions above. However, we can make a few simplifying assumptions. First, as mentioned above, the distribution of particle arrival rate $P\left(\{r(t)\} \mid s(t+\tau)\right)$ has Poisson statistics, but if a sufficient number of particles arrive at the cell's receptor array per unit time, the Poisson statistics are approximately Gaussian.

Even with this approximation, $P(\{r(t)\}) = \int P\left(\{r(t)\} \mid s(t+\tau)\right) P\left(s(t+\tau)\right) ds$ is not Gaussian because $P\left(s(t+\tau)\right)$ is not Gaussian. However, in shallow gradients (small $s$), the (roughly) Gaussian particle arrival noise described by $P\left(\{r(t)\} \mid s(t+\tau)\right)$ blurs the non-Gaussian structure in $P\left(s(t+\tau)\right)$, making $P(\{r(t)\})$ nearly Gaussian, as well. As a result, we can approximate the mutual information in Eqn. 22 by approximating all distributions as Gaussian, as shown rigorously by others (12–14).

Since all distributions are approximately Gaussian, the posterior distribution of $s(t+\tau)$ given past $r(t)$ is Gaussian as well, with mean $\hat{s}_r(t+\tau)$ and variance $\sigma^2_{s|r}(\tau)$ : $P(s(t+\tau)|\{r(t)\}) = \mathcal{N}\left(\hat{s}_r(t+\tau), \sigma^2_{s|r}(\tau)\right)$. With this, the mutual information can be computed from:

$$I\left(\{r(t)\}; s(t+\tau)\right) \approx \frac{1}{2}\log\left(\frac{\sigma_s^2}{\sigma^2_{s|r}(\tau)}\right) = -\frac{1}{2}\log\left(1-\rho^2_{rs}(\tau)\right), \quad (23)$$

and the information rate is:

$$\dot{I}^*_{s\to r} = \frac{1}{2}\left[\frac{-\partial_\tau \rho^2_{rs}(\tau)}{1-\rho^2_{rs}(\tau)}\right]_{\tau=0}. \quad (24)$$

Above, $\sigma_s^2 = g^2\,\sigma_v^2$ is the marginal variance of $s(t+\tau)$ or $s(t)$, i.e. the variance of the distribution $P\left(s(t+\tau)\right) = P\left(s(t)\right)$ (by time-translation invariance). Then, $\rho^2_{rs}(\tau) = 1 - \frac{\sigma^2_{s|r}(\tau)}{\sigma_s^2}$ is a generalized correlation coefficient between $s(t+\tau)$ and past $r$, or the fraction reduction of variance in $s(t+\tau)$ upon observing past $r$.

To calculate $\dot{I}^*_{s\to r}$, we now need to calculate the generalized correlation coefficient $\rho^2_{rs}(\tau)$, or the posterior variance of $s(t+\tau)$, $\sigma^2_{s|r}(\tau)$, using the models for the dynamics of $s$ and $r$ above. To do this, we first note that the posterior mean of $s(t+\tau)$, $\hat{s}_r(t+\tau)$, can be computed using the causal Wiener filter, $M_r(T)$, which minimizes the following mean squared error $\langle e^2(\tau)\rangle$:

$$\langle e^2(\tau)\rangle = \left\langle \left( s(t+\tau) - \int_{-\infty}^{t} M_r(t-t')\, r(t')\, dt' \right)^2 \right\rangle \tag{25}$$

Once the optimal kernel $M_r(T)$ is obtained, the posterior mean is $\hat{s}_r(t+\tau) = \int_{-\infty}^{t} M_r(t-t')\, r(t')\, dt'$ and the posterior variance is $\sigma^2_{s|r}(\tau) = \langle e^2(\tau)\rangle$ . Therefore, to derive the mutual information $I\big(\{r(t)\}; s(t+\tau)\big)$, and thus the information rate $\dot{I}^*_{s\to r}$, we need to derive this Wiener filter. The main challenge in deriving $M_r(T)$ is that it must satisfy the constraint that it is causal: that is, we require that $M_r(T) = 0$ for $T < 0$. In Appendix A, we derive the necessary equations and explain where they come from, but here we will just apply them to get $M_r(t)$. See also references (4,15,16).

The optimal kernel can be expressed in Fourier space terms of the power spectra of the signal $s(t)$ and the particle arrival rate $r(t)$ as (Appendix A):

$$M_r(\omega) = \frac{1}{\phi_r(\omega)} \left[ \frac{S_{rs}(\omega)}{\phi_r^*(\omega)}\, e^{-i\,\omega\,\tau} \right]^+ . \tag{26}$$

$M_r(\omega)$ is the Fourier transform of $M_r(T)$. $\phi_r(\omega)$ is the causal part of the spectral decomposition of $S_r(\omega)$ (defined below and in Appendix A), where $S_r(\omega)$ is the power spectrum of $r$. $\phi_r^*(\omega)$ is its (anti-causal) complex conjugate. $S_{rs}(\omega)$ is the cross-spectra of $r$ and $s$, equivalent to the Fourier transform of $C_{rs}(\tau)$, where $C_{rs}(\tau) = \langle (r(t) - r_0) s(t+\tau)\rangle$ is the cross-correlation of $s$ and $r$ in the time domain. Finally, $[f(\omega)]^+$ indicates the causal part of the inverse Fourier transform of $f(\omega)$, which can be found by taking the inverse Fourier transform of $f(\omega)$, multiplying the result by a Heaviside step function in the time domain, and then taking the Fourier transform.

As explained in Appendix A, to find the optimal causal kernel, we need to decompose $S_r(\omega)$ into the product of a causal and an anti-causal part. This requires finding the zeros and poles of $S_r(\omega)$. The zeros satisfy $S_r(\omega = i\, z_r) = 0$, and therefore are the complex solutions to the equation:

$$2\, r_0\, g^2\, \sigma_v^2\, \tau_v^3 + (\epsilon^2 + \tau_v^2\, \omega^2)\,(1 + \tau_v^2\, \omega^2) = 0 \tag{27}$$

or, defining $\gamma_r = 2\, r_0\, g^2\, \sigma_v^2\, \tau_v^3$:

$$\gamma_r + (\epsilon^2 + \tau_v^2\, \omega^2)\,(1 + \tau_v^2\, \omega^2) = 0. \tag{28}$$

$\gamma_r$ is a dimensionless signal-to-noise ratio parameter, where the signal is $r_0^2\, g^2\, \sigma_v^2\, \tau_v^3$ (the prefactor of the first term in $S_r(\omega)$ when $\omega$ is rescaled by $1/\tau_v$) and the noise is $r_0$ (the second term in $S_r(\omega)$).

The zeros of $S_r(\omega)$ are:

$$i\, z_{r,1} = i \frac{1}{\sqrt{2}\,\tau_v} \sqrt{1 + \epsilon^2 + \sqrt{(1-\epsilon^2)^2 - 4\,\gamma_r}}, \qquad i\, z_{r,2} = i \frac{1}{\sqrt{2}\,\tau_v} \sqrt{1 + \epsilon^2 - \sqrt{(1-\epsilon^2)^2 - 4\,\gamma_r}}, \tag{29}$$

as well as their complex conjugates, $z_1^*$ and $z_2^*$. As $\epsilon \to 0$, these will simplify to:

$$i\, z_{r,1} = i\, \frac{1}{\sqrt{2}\,\tau_v} \sqrt{1 + \sqrt{1 - 4\,\gamma_r}}, \qquad i\, z_{r,2} = i\, \frac{1}{\sqrt{2}\,\tau_v} \sqrt{1 - \sqrt{1 - 4\,\gamma_r}}, \tag{30}$$

Note that there are several equivalent forms for these zeros, and they change from being fully imaginary to complex when $\gamma_r > 1/4$.

The poles of $S_r(\omega)$ satisfy $\frac{1}{S_r(\omega = i\, p_r)} = 0$ and are given by $i\, p_{r,1} = i \frac{\epsilon}{\tau_v}$ and $i\, p_{r,2} = i \frac{1}{\tau_v}$, as well as their complex conjugates $p_{r,1}^*$ and $p_{r,2}^*$.

Power spectral densities of real, stable, causal systems can generally be decomposed into causal and anti-causal parts ("Wiener-Hopf factorization") (17–19):

$$S_r(\omega) = \phi_r(\omega)\, \phi_r^*(\omega) \tag{31}$$

where

$$\phi_r(\omega) = \sqrt{r_0}\, \frac{\left(z_{r,1} - i\,\omega\right)\left(z_{r,2} - i\,\omega\right)}{\left(p_{r,1} - i\,\omega\right)\left(p_{r,2} - i\,\omega\right)} \tag{32}$$

has zeros and poles with negative imaginary parts, and $\phi_r^*(\omega)$ is its complex conjugate.

Next, we need the causal part of (see Appendix A):

$$\frac{S_{rs}(\omega)}{\phi_r^*(\omega)} e^{-i\,\omega\,\tau} = \frac{\sqrt{r_0}}{\frac{\epsilon}{\tau_v} + i\,\omega} S_s(\omega) \frac{\left(p_{r,1} + i\,\omega\right)\left(p_{r,2} + i\,\omega\right)}{\left(z_{r,1} + i\,\omega\right)\left(z_{r,2} + i\,\omega\right)} e^{-i\,\omega\,\tau} \tag{33}$$

$$= \frac{\sqrt{r_0}}{\frac{\epsilon}{\tau_v} + i\,\omega} \frac{2\, g^2 \frac{\sigma_v^2}{\tau_v}}{\frac{1}{\tau_v^2} + \omega^2} \frac{\left(\frac{\epsilon}{\tau_v} + i\,\omega\right)\left(\frac{1}{\tau_v} + i\,\omega\right)}{\left(z_{r,1} + i\,\omega\right)\left(z_{r,2} + i\,\omega\right)} e^{-i\,\omega\,\tau} \tag{34}$$

$$= \sqrt{r_0}\, \frac{2\, g^2 \frac{\sigma_v^2}{\tau_v}}{\left(\frac{1}{\tau_v} - i\,\omega\right)} \frac{1}{\left(z_{r,1} + i\,\omega\right)\left(z_{r,2} + i\,\omega\right)} e^{-i\,\omega\,\tau} \tag{35}$$

$$= \frac{\gamma_r}{\sqrt{r_0}\, \tau_v^4} \frac{1}{\left(\frac{1}{\tau_v} - i\,\omega\right)} \frac{1}{\left(z_{r,1} + i\,\omega\right)\left(z_{r,2} + i\,\omega\right)} e^{-i\,\omega\,\tau} \tag{36}$$

One approach would be to compute the inverse Fourier transform of $\frac{S_{rs}(\omega)}{\phi_r^*(\omega)}$, apply the time shift forward by $\tau$ implied by $e^{-i\,\omega\,\tau}$, multiply the result by a Heaviside step function $\Theta(T)$, and compute the Fourier transform of the result. An alternative approach is to compute the partial fraction decomposition of the expression above and keep only the terms with poles and zeros that have negative imaginary part:

$$\frac{S_{rs}(\omega)}{\phi_r^*(\omega)} e^{-i\,\omega\,\tau} = \frac{A}{\left(\frac{1}{\tau_v} - i\,\omega\right)} + \frac{B}{\left(z_{r,1} + i\,\omega\right)} + \frac{C}{\left(z_{r,2} + i\,\omega\right)} \tag{37}$$

for unknown $A$, $B$, and $C$. Only the pole of the first term ($\omega = -i \frac{1}{\tau_v}$) has negative imaginary part, so we only need to compute $A$ to get the causal part of this expression. With some algebra, this is:

$$A = \left[\frac{\gamma_r}{\sqrt{r_0}\,\tau_v^4}\frac{1}{\left(z_{r,1} + i\,\omega\right)\left(z_{r,2} + i\,\omega\right)}e^{-i\,\omega\,\tau}\right]_{\omega=-i\frac{1}{\tau_v}} \tag{38}$$

$$= \frac{\gamma_r}{\sqrt{r_0}\,\tau_v^2}\frac{1}{\left(1 + \tau_v\,z_{r,1}\right)\left(1 + \tau_v\,z_{r,2}\right)}e^{-\frac{\tau}{\tau_v}}, \tag{39}$$

and the causal part of $\frac{S_{rs}(\omega)}{\phi_r^*(\omega)}e^{-i\,\omega\,\tau}$ is then:

$$\left[\frac{S_{rs}(\omega)}{\phi_r^*(\omega)}e^{-i\,\omega\,\tau}\right]^+ = \frac{\gamma_r}{\sqrt{r_0}\,\tau_v^2}\frac{1}{\left(1 + \tau_v\,z_{r,1}\right)\left(1 + \tau_v\,z_{r,2}\right)}\frac{1}{\left(\frac{1}{\tau_v} - i\,\omega\right)}e^{-\frac{\tau}{\tau_v}}. \tag{40}$$

Finally, the optimal kernel that computes the mean of $p(s(t+\tau)|\{r(t)\})$, $\mu_{s|r}(\tau)$, is (Appendix A):

$$M_r(\omega) = \frac{1}{\phi_r(\omega)}\left[\frac{S_{rs}(\omega)}{\phi_r^*(\omega)}e^{-i\,\omega\,\tau}\right]^+ \tag{41}$$

$$= e^{-\frac{\tau}{\tau_v}}\,\frac{\gamma_r}{r_0\,\tau_v^2}\frac{1}{\left(1 + \tau_v\,z_{r,1}\right)\left(1 + \tau_v\,z_{r,2}\right)}\frac{-i\,\omega}{\left(z_{r,1} - i\,\omega\right)\left(z_{r,2} - i\omega\right)}, \tag{42}$$

after taking $\epsilon$ to zero. We convert this kernel to the time domain and discuss its properties in the next section.

The variance of $p(s(t+\tau)|\{r(t)\})$, $\sigma_{s|r}^2(\tau)$, is (Appendix A, Eqn. 181):

$$\sigma_{s|r}^2(\tau) = \sigma_s^2 - \frac{1}{2\pi}\int_{-\infty}^{\infty} S_{rs}^*(\omega)\,e^{i\,\omega\,\tau}\,M_r(\omega)\,d\omega \tag{43}$$

$$= \sigma_s^2\left(1 - e^{-2\frac{\tau}{\tau_v}}\frac{\gamma_r}{\left(1 + \tau_v\,z_{r,1}\right)^2\left(1 + \tau_v\,z_{r,2}\right)^2}\right) \tag{44}$$

where we used $\sigma_s^2 = g^2\sigma_v^2 = \gamma/(2\,r_0\,\tau_v^3)$. Then the correlation coefficient $\rho_{rs}^2(\tau)$ is:

$$\rho_{rs}^2(\tau) = 1 - \frac{\sigma_{s|r}^2(\tau)}{\sigma_s^2} = e^{-2\frac{\tau}{\tau_v}}\frac{\gamma_r}{\left(1 + \tau_v\,z_{r,1}\right)^2\left(1 + \tau_v\,z_{r,2}\right)^2}. \tag{45}$$

Finally, using Eqn. 24 from above, we find that the behaviorally-relevant information available in particle counts is:

$$\dot{I}_{s\to r}^* = \frac{1}{\tau_v}\frac{\rho_{rs}^2(\tau=0)}{1 - \rho_{rs}^2(\tau=0)} = \frac{1}{\tau_v}\,\frac{\gamma_r\frac{1}{\left(1 + \frac{1}{\sqrt{2}}\sqrt{1 + \sqrt{1 - 4\,\gamma_r}}\right)^2\left(1 + \frac{1}{\sqrt{2}}\sqrt{1 - \sqrt{1 - 4\,\gamma_r}}\right)^2}}{1 - \gamma_r\frac{1}{\left(1 + \frac{1}{\sqrt{2}}\sqrt{1 + \sqrt{1 - 4\,\gamma_r}}\right)^2\left(1 + \frac{1}{\sqrt{2}}\sqrt{1 - \sqrt{1 - 4\,\gamma_r}}\right)^2}}. \tag{46}$$

Expanding around small SNR, $\gamma_r$, gives:

$$\dot{I}^*_{s\to r} \approx \frac{1}{\tau_v}\frac{\gamma_r}{4} = \frac{1}{2} r_0\, g^2\, \sigma_v^2\, \tau_v^2. \tag{47}$$

Note that for small signals, Eqn. 46 can be written $\dot{I}^*_{s\to r} \approx \frac{1}{\tau_v}\rho_{rs}^2(\tau = 0) \approx \frac{2}{\tau_v} I\big(\{r(t)\}; s(t)\big)$.

The optimal kernel in the time domain and the information rate remain real when $\gamma_r > 1/4$, even though $z_{r,1}$ and $z_{r,2}$ become complex. In this regime, they can be written:

$$z_{r,1} = \frac{1}{2\,\tau_v}\left(\sqrt{2\sqrt{\gamma_r}+1} - i\sqrt{2\sqrt{\gamma_r}-1}\right), \qquad z_{r,2} = \frac{1}{2\,\tau_v}\left(\sqrt{2\sqrt{\gamma_r}+1} + i\sqrt{2\sqrt{\gamma_r}-1}\right). \tag{48}$$

The optimal kernel in frequency space can be written:

$$M_r(\omega) = e^{-\frac{\tau}{\tau_v}}\, \frac{\gamma_r}{r_0\,\tau_v^2}\frac{1}{\left(1+\tau_v\, z_{r,1}\right)\left(1+\tau_v\, z_{r,2}\right)}\frac{-i\,\omega}{\left(z_{r,1} - i\,\omega\right)\left(z_{r,2} - i\omega\right)} \tag{49}$$

$$= e^{-\frac{\tau}{\tau_v}}\, \frac{\gamma_r}{r_0\,\tau_v^2}\frac{1}{\left(1+\sqrt{1+2\sqrt{\gamma_r}}+\sqrt{\gamma_r}\right)}\frac{-i\,\omega}{\left(z_{r,1} - i\,\omega\right)\left(z_{r,2} - i\omega\right)}, \tag{50}$$

the correlation coefficient at $\tau = 0$ can be written:

$$\rho_{rs}^2(\tau = 0) = \frac{\gamma_r}{\left(1+z_{r,1}\,\tau_v\right)^2\left(1+z_{r,2}\,\tau_v\right)^2} \tag{51}$$

$$= \frac{\gamma_r}{\left|1+z_{r,1}\,\tau_v\right|^4} \tag{52}$$

$$= \frac{\gamma_r}{\left(1+\sqrt{1+2\sqrt{\gamma_r}}+\sqrt{\gamma_r}\right)^2}, \tag{53}$$

and the information rate is:

$$\dot{I}^*_{s\to r} = \frac{1}{\tau_v}\frac{\dfrac{\gamma_r}{\left(1+\sqrt{1+2\sqrt{\gamma_r}}+\sqrt{\gamma_r}\right)^2}}{1-\dfrac{\gamma_r}{\left(1+\sqrt{1+2\sqrt{\gamma_r}}+\sqrt{\gamma_r}\right)^2}}. \tag{54}$$

For small $\gamma_r$, this reduces again to Eqn. 47.

#### Correlation coefficients

As described above, in shallow gradients, the posterior distribution $P(s(t)|\{r\})$ is approximately Gaussian with constant variance, $\sigma^2_{s(t)|r}$, and time-varying mean, $\hat{s}_r(t)$, that depends on $\{r\}$. Once $\hat{s}_r(t)$ is computed, the posterior $P(s(t)|\{r\}) = P\big(s(t)\big|\hat{s}_r(t)\big)$ does not depend on $\{r\}$ in any other way, so $\hat{s}_r(t)$ is a sufficient statistic. This in turn means that the posterior variance of $s(t)$ given $\hat{s}_r(t)$ is the same as the posterior variance of $s(t)$ given $\{r\}$, and thus the reduction in uncertainty about $s(t)$ by observing $\{r\}$ or $\hat{s}_r(t)$ is the same. Since the generalized correlation coefficient $\rho_{rs}^2$ quantifies this reduction in uncertainty about $s(t)$ by observing $\{r\}$, $\rho_{rs}^2$ must also be the squared correlation between $s(t)$ and $\hat{s}_r(t)$. The same applies if we replace $r$ with $a$.

#### Optimal kernel for estimating signal from particle arrivals

To get the time-domain kernel mapping past particle arrival rate $r(t)$ to signal $s(t+\tau)$, $M_r(T)$,we take the inverse Fourier transform of $M_r(\omega)$. $M_r(T)$ has the form of a sum of two exponentials, with real exponents when $\gamma_r \leq 1/4$ and complex ones when $\gamma_r > 1/4$. For $\gamma_r < 1/4$, the kernel in the time domain is:

$$M_r(T) = e^{-\frac{\tau}{\tau_v}} \frac{\gamma_r}{r_0\, \tau_v^2} \frac{1}{\left(1+z_{r,1}\,\tau_v\right)\left(1+z_{r,2}\,\tau_v\right)} \frac{\left(z_{r,1}\, e^{-z_{r,1}\,T} - z_{r,2}\, e^{-z_{r,2}\,T}\right)}{z_{r,1}-z_{r,2}}\, \Theta(T), \tag{55}$$

where $\Theta(T)$ is the Heaviside step function, indicating that the kernel is indeed causal.

The optimal kernel $M_r(T)$ essentially computes the time derivative of concentration, while also averaging out shot noise from particle arrivals. It has several notable features. First, it is biphasic and exhibits perfect adaptation, a hallmark of the chemotaxis pathway. Any derivative operation should adapt perfectly because it should only respond to *changes* in the input.

It is interesting to examine how the time scales of the optimal kernel are set by the signal-to-noise ratio $\gamma_r = 2\, r_0\, g^2\, \sigma_v^2\, \tau_v^3$. The initial response time scale is set by $z_{r,1}^{-1}$ and its adaptation time scale is set by $z_{r,2}^{-1}$. When the inputs are very noisy, i.e. as $\gamma_r \to 0$, $z_{r,1}^{-1}$ gets longer but saturates at $\tau_v$:

$$z_{r,1}^{-1}(\gamma_r \to 0) \;\approx\; \tau_v\left(1-\frac{\gamma_r}{2}\right). \tag{56}$$

This makes sense because it maximally averages out shot noise, but only for as long as past signals are correlated with the current signal. As the SNR increases, this initial averaging time gets shorter.

As the inputs get noisier, i.e. as $\gamma_r \to 0$, the adaptation time approaches:

$$z_{r,2}^{-1}(\gamma_r \to 0) \;\approx\; \tau_v\left(\frac{1}{\sqrt{\gamma_r}} - \frac{\sqrt{\gamma_r}}{2}\right). \tag{57}$$

This shows that the adaptation time can become long compared to $\tau_v$ when $\gamma_r < 1/4$.

Interestingly, in this regime, the kernel $M_r(T)$ has the same functional form as the phenomenological kernel we measured previously (1) (after transforming the input quantity from $s(t)$ to $c(t)$).

When signal and noise have similar strength $\gamma_r = 1/4$, $z_{r,1} = z_{r,2} = z_r = \frac{1}{\sqrt{2}}\,\tau_v^{-1}$, and the optimal kernel becomes:

$$M_r(T) = e^{-\frac{\tau}{\tau_v}} \frac{\gamma_r}{r_0\,\tau_v^2} \frac{1}{\left(1+\frac{1}{\sqrt{2}}\right)^2} e^{-z_r\,T}(1 - z_r\,T)\,\Theta(T). \tag{58}$$

When SNR is high $\gamma_r > 1/4$, $z_{r,1}$ and $z_{r,2}$ become complex. However, since they are complex conjugates of each other, the kernel remains real:

$$M_r(T) = e^{-\frac{\tau}{\tau_v}}\,\frac{\gamma_r}{r_0\,\tau_v^2}\frac{1}{\left(1+z_{r,1}\,\tau_v\right)\left(1+z_{r,2}\,\tau_v\right)}\,e^{-\mathrm{Re}[z_{r,2}]\,T}\left(\cos\left(\mathrm{Im}\left[z_{r,2}\right]T\right)-\frac{\mathrm{Re}\left[z_{r,2}\right]}{\mathrm{Im}\left[z_{r,2}\right]}\sin\left(\mathrm{Im}\left[z_{r,2}\right]T\right)\right)\Theta(T)$$

$$= e^{-\frac{\tau}{\tau_v}}\,\frac{\gamma_r}{r_0\,\tau_v^2}\frac{1}{\left(1+\sqrt{1+2\sqrt{\gamma_r}}+\sqrt{\gamma_r}\right)}\,e^{-\frac{1}{2}\sqrt{2\sqrt{\gamma_r}+1}\frac{T}{\tau_v}}\times$$

$$\left(\cos\left(\frac{1}{2}\sqrt{2\sqrt{\gamma_r}-1}\,\frac{T}{\tau_v}\right)-\sqrt{\frac{2\sqrt{\gamma_r}+1}{2\sqrt{\gamma_r}-1}}\sin\left(\frac{1}{2}\sqrt{2\sqrt{\gamma_r}-1}\,\frac{T}{\tau_v}\right)\right)\Theta(T) \qquad (59)$$

The optimal kernel, $M_r(T)$, is plotted in Fig. S4 for varying values of $\gamma_r$.

As the SNR $\gamma_r$ increases, the initial response time and the adaptation time both get shorter. Although the kernel oscillates, its decay rate is faster than the period of oscillations. The time scales of decay and oscillation are closest to each other, and thus the oscillation amplitude is largest, when $\gamma_r$ is large: in the limit that $\gamma_r \to \infty$, $\mathrm{Re}\left[z_{r,2}\right] = \mathrm{Im}\left[z_{r,2}\right] = {\gamma_r}^{1/4}$. Even in this limit, the peak of the kernel following the first negative lobe occurs at time $T = 3\pi\,{\gamma_r}^{-1/4}$ and is smaller than the kernel's maximum value ($M_r(T=0)$) by a factor of $e^{-3\pi/2} \sim 0.009$. Thus, the oscillations are small. $M_r(T)$ transitions continuously between the forms above as $\gamma_r$ varies.

The results of this and previous section could also be derived using the continuous-time Kalman-Bucy filter (20,21). That approach provides a pair of ODEs for the estimator of $s$ (i.e. conditional mean $\mu_{s|r}$) and its uncertainty (i.e. the conditional variance $\sigma_{s|r}^2$) that are driven by the observations, $r(t)$. Once $\sigma_{s|r}^2$ reaches steady state in that formulation (consistent with our assumption of stationarity here), the ODE for $\mu_{s|r}$ can be solved in terms of a kernel convolved with past $r(t)$, which is identical to the optimal kernel above.

#### Comparing temporal and spatial sensing

Berg and Purcell argued that bacteria are too small to accurately infer concentration differences across their body. We can estimate the effect of measuring concentration differences across the cell body by comparing its SNR to that of temporal comparisons. For temporal comparisons, we showed above that the SNR is $\gamma_r = 2\,r_0\,g^2\,\sigma_v^2\,\tau_v^3 = 2\,(r_0\,\tau_v)\,(g^2\,\sigma_v^2\,\tau_v^2) = 2\,(r_0\,\tau_v)\,\left(L_{run}/L_{gradient}\right)^2$. Here, we defined the run length scale $L_{run} \approx \sigma_v\,\tau_v$ and the gradient length scale $L_{gradient} = g^{-1}$. When making spatial comparisons instead of temporal ones, the run length is replaced by the length over which spatial comparisons are made, i.e. the cell body length, $L_{body}$: $\gamma_r^{spatial} = 2\,(r_0\,\tau_v)\,\left(L_{body}/L_{gradient}\right)^2$. The ratio of these two SNR's is $\gamma_r^{spatial}/\gamma_r = \left(L_{body}/L_{run}\right)^2$. The length of the cell is roughly 2 μm and its run length is roughly 20 μm, making $\gamma_r^{spatial}/\gamma_r \approx 1/100$. Therefore, consistent with Berg and Percell's arguments, spatial sensing has a negligible effect on sensing accuracy for bacteria with run lengths much longer than the body length.

#### Berg & Purcell's SNR threshold for chemotaxis

Berg and Purcell claimed that *E. coli* chemosensing approaches the physical limit (7), based on the following argument. They postulated that, in order to perform chemotaxis, a cell must estimate the change in concentration over a single run with uncertainty smaller than the concentration change itself.

They envisioned the cell making two measurements of concentration, $c$, during a run, each of duration $T$. Their condition for chemotaxis to be possible, with a slight change in notation, reads (Eqn. 57 in Ref. (7)):

$$\tau_v \frac{dc}{dt} > \sqrt{2}\, \delta c(T). \tag{60}$$

The left-hand side is the concentration change over a single run of duration $\tau_v$, and the right-hand side is the standard deviation of the estimated concentration change (i.e. the uncertainty of the difference of two independent measurements of concentration, $c_1$ and $c_2$, is $\sqrt{\delta c_1^2 + \delta c_2^2} \approx \sqrt{2}\, \delta c$), which depends on the measurement time $T$. Equating the two sides above sets a minimum time for each measurement, $T^{min}$, and the argument is that chemotaxis would not be possible if the cell's run duration is shorter than the time needed to make the two measurements. Thus, chemotaxis requires $\tau_v > 2\, T^{min}$. Conversely, if a cell's run duration is much longer than $2\, T^{min}$, that would imply that the cell's sensing accuracy is far below the physical limit. Thus, they argue that an ideal agent would exhibit an optimal run duration of:

$$\tau_v^* \approx 2\, T^{min}. \tag{61}$$

Plugging realistic parameters, they argued that this equality approximately holds. Based on this, they concluded, that "[t]he chemotactic sensitivity of *Escherichia coli* approaches that of the cell of optimal design."

The problem with their argument is the initial postulate: to perform chemotaxis, *E. coli* do not have to detect concentration changes with significance above a strict threshold *in a single run*. Instead, they only need to increase their run duration when the concentration increases, and decrease it when concentration decreases, *on average*. This is because chemotaxis drift speed depends on an *average* over many runs. This is consistent with our previous result (1) that the total amount of information that *E. coli* acquire in a single run ($\sim 1$ second) is much smaller than 1 bit, the amount required to unambiguously distinguish whether concentration increased or decreased over a single run.

Next, we provide a counterexample to the argument above using simulations, showing that chemotaxis is possible even when Eqn. 60 is not satisfied. First, we express their condition in terms of our signal-to-noise ratio of particle arrivals, $\gamma_r$. For an ideal agent with a circular sensor of radius $l$ that uptakes arrival particles, the particle arrival flux is $J = r_0$ and the average receptor occupancy is $\bar{p} = 0$ because the receptors are always free to sense particle arrivals. Then Berg and Purcell showed that the ideal agent's uncertainty about a measurement of concentration is (Eqn. 55 of (7)):

$$\delta c = \frac{c_0}{\sqrt{\frac{1}{2}\, r_0\, T}}. \tag{62}$$

If the cell makes two consecutive measurements in a single run, each of duration $T = \frac{\tau_v}{2}$, the condition for chemotaxis in Eq. 60 becomes:

$$\tau_v \frac{dc}{dt} > 2\sqrt{2} \frac{c_0}{\sqrt{r_0 \tau_v}}. \tag{63}$$

In a single up-gradient run, $\frac{dc}{dt} \approx v_0 \frac{dc}{dx}$, and the equation above can be written as:

$$\tau_v\, v_0\, \frac{dc}{dx} > \; 2\sqrt{2}\frac{c_0}{\sqrt{r_0 \tau_v}}. \tag{64}$$

Then, since $\frac{dc}{dx} \approx c_0 \frac{d}{dx} \log(c) = c_0\, g$, we get:

$$\tau_v\, v_0\, g > \frac{2\sqrt{2}}{\sqrt{r_0 \tau_v}} \tag{65}$$

$$r_0\, v_0^2\, g^2\, \tau_v^3 > 8 \tag{66}$$

$$2\, r_0 \frac{v_0^2}{3} g^2\, \tau_v^3 > \frac{16}{3} \tag{67}$$

$$2\, r_0\, \sigma_v^2\, g^2\, \tau_v^3 > \frac{16}{3} \tag{68}$$

$$\gamma_r > \frac{16}{3}. \tag{69}$$

Eqn. 68 used $\sigma_v^2 = \frac{1}{3} v_0^2$ when tumbles are instantaneous. Thus, Berg and Purcell's condition for chemotaxis to be possible in Eqn. 60 is equivalent to requiring that $\gamma_r > 16/3$. However, we show in simulations (Fig. 4B of the main text) that ideal cells, and even *E. coli*, can clearly climb the gradient when $g = 0.05\ \mathrm{mm}^{-1}$, $c_0 = 1\ \mu\mathrm{M}$, and $\gamma_r = 0.015 \ll 16/3$.

#### Modeling kinase activity

In shallow gradients, CheA kinases respond approximately linearly to recent signals. We model kinase responses, $a(t)$, to past particle arrival rates, $r(t)$, in background particle arrival rate $r_0$, as:

$$a(t) = a_0 - \int_{-\infty}^{t} K_r(t - t')\, (r(t') - r_0)\, dt' + \eta(t). \tag{70}$$

The response function to particle arrival rate, $K_r(T)$, is:

$$K_r(T) = G_r \left( \left(\frac{1}{\tau_1} + \frac{1}{\tau_2}\right) \exp\left(-\left(\frac{1}{\tau_1} + \frac{1}{\tau_2}\right) T\right) - \frac{1}{\tau_2} \exp\left(-\frac{T}{\tau_2}\right) \right) \Theta(T), \tag{71}$$

where $\tau_1$ and $\tau_2$ above have the same meaning as in our previous work (1). In the main text, we replaced $\left(\frac{1}{\tau_1} + \frac{1}{\tau_2}\right) \to \frac{1}{\tau_1}$ for space, since $\tau_2 \gg \tau_1$.

The Fourier transform of this kernel is:

$$K_r(\omega) = \frac{G_r}{\tau_1} \frac{(-i\, \omega)}{\left(\frac{1}{\tau_2} - i\, \omega\right)\left(\frac{1}{\tau_1} + \frac{1}{\tau_2} - i\, \omega\right)}. \tag{72}$$

In our previous work (1), we modeled responses of kinase activity to past signals $s$ instead of past particle arrival rate $r$. These two descriptions are equivalent in the regime of shallow gradients. We show this below by starting from average responses of kinase activity to particle arrival rate:

$$\langle a(t)\rangle = a_0 - \int_{-\infty}^{t} K_r(t-t')\,(\langle r(t')\rangle - r_0)\,dt' \,, \tag{73}$$

where angled brackets indicate averaging over repeated presentation of the same signal trajectory $\{s\}$, and thus they average out particle noise and kinase noise. From here, we will derive a response kernel to past signals that gives identical kinase responses.

First, we note that:

$$\langle r(t)\rangle - r_0 = k_D\,(c(t) - c_0) = r_0 \int_{-\infty}^{t} s(t')\,dt' \,, \tag{74}$$

where we used $s(t) \approx \frac{1}{c_0}\frac{dc}{dt}$ in shallow gradients. Then, it is convenient to transform the expressions above to Fourier space, where $\delta a(\omega) = F[\langle a(t)\rangle - a_0]$ , $\delta r(\omega) = F[\langle r(t)\rangle - r_0]$ , and $K_r(\omega) = F[K_r(T)]$, and again $F[f(t)] = \int_{-\infty}^{\infty} f(t)\, e^{i\,\omega\,t}\, dt$ is the Fourier transform. Then we have:

$$\delta a(\omega) = -K_r(\omega)\,\delta r(\omega), \tag{75}$$

$$\delta r(\omega) = r_0 \frac{s(\omega)}{-i\,\omega}. \tag{76}$$

With this, we get:

$$\delta a(\omega) = -K_r(\omega)\, r_0 \frac{s(\omega)}{-i\,\omega} = -K(\omega)\, s(\omega) \tag{77}$$

where $K(\omega) = r_0 \frac{K_r(\omega)}{-i\,\omega}$ is the Fourier transform of the linear response function to signals. Thus, we can either write down average kinase responses to particle arrival rate $r(t)$, with linear response function $K_r(T)$, or responses to signals $s(t)$, with linear response function $K(T)$ (1):

$$K(T) = r_0 \int_0^T K_r(t')\,dt' = G \exp\left(-\frac{T}{\tau_2}\right)\left(1 - \exp\left(-\frac{T}{\tau_1}\right)\right)\Theta(T). \tag{78}$$

Here, we have defined the MWC model gain $G = r_0\, G_r$ (22,23). In the MWC model, kinase-receptor complexes can be in active or inactive states. The dissociation constants for the attractant in each state, $K_i$ and $K_a$, are different, with $K_i \ll K_a$, which causes attractant concentration to influence the fraction of kinases in the active state. When the background concentration $c_0 \ll K_a$, the gain of the kinase response to changes in log-concentration of attractant can be written:

$$G(c_0) \approx G_\infty \frac{c_0}{c_0 + K_i}. \tag{79}$$

where $G_\infty$ is the “log-sensing” gain (when $c_0 \gg K_i$).

We can use the response function to particle arrivals, $K_r(T)$, to compute the power spectrum of particle counting noise filtered through the kinase response kernel, $K_r(T)$, but expressed it in terms of the response kernel $K(T)$ to signals $s$. Since we model particle arrival noise as shot noise, its power spectrum is constant and equal to $r_0$. Filtering this noise through the response kernel $K_r(\omega)$ gives:

$$N_r(\omega) = r_0\,|K_r(\omega)|^2 = r_0\,\left|\frac{-\,i\,\omega}{r_0}\,K(\omega)\right|^2 = \frac{1}{r_0}\,\omega^2\,|K(\omega)|^2. \tag{80}$$

In experiments, we measure responses to absolute changes in concentration $c(t)$, with response kernel $K_c(t)$, which has the same form as $K_r(T)$ above, but with gain $G_c$. Then, we convert $G_c$ to $G_r$ via $G_r = G_c/k_D$, and thus convert $K_c(T)$ to $K_r(T)$. With this, the intensity of filtered particle noise in Eqn. 80 is proportional to $G_r^2\,r_0 = G_c^2\,c_0/k_D$. This conversion implies that *E. coli* respond to every particle arriving at their receptors, which is unlikely. Instead, one might use an effective $k_D^{eff} < k_D$ to do the conversion above, which would increase our estimate for the intensity of filtered particle noise, being proportional to $G_c^2\,c_0/k_D^{eff}$. However, modeling the filtered particle noise with $k_D^{eff} = k_D$ maximizes our estimate of *E. coli*'s information rate. Since we find that *E. coli* are far from the physical limit, this is a conservative modeling choice. Furthermore, we believe $k_D^{eff}$ can't be much smaller than $k_D$, otherwise kinase activity noise would exhibit significant concentration-dependence, which we do not observe experimentally (SI Fig. S1C).

Next, we consider modeling noise in kinase activity. As explained in the main text and in Fig. S3, the FRET system we use for measuring kinase activity has limited time resolution, about $0.3\ s$. This allows us to constrain slow fluctuations in kinase activity, whose correlation function is characterized by a single decaying exponential function (1,24):

$$\langle \eta(t)\eta(t')\rangle = N_n(t-t') = \sigma_n^2 \exp\left(-\frac{|t-t'|}{\tau_n}\right) = D_n\,\tau_n \exp\left(-\frac{|t-t'|}{\tau_n}\right). \tag{81}$$

The parameters here are the long-time variance $\sigma_n^2$ and the correlation time $\tau_n$, which are related to the diffusivity of the noise by $D_n = \sigma_n^2/\tau_n$. The power spectrum of this noise is

$$N_n(\omega) = \frac{2\,D_n}{\frac{1}{\tau_n^2} + \omega^2}. \tag{82}$$

There can also be noise at higher frequencies that we don't observe. Kinase responses to particle arrival noise set a minimum noise level at all frequencies. At high frequencies, simply extrapolating the power spectrum in Eqn. 82 drops below the implied filtered particle noise in Eqn. 80 if we take $\tau_1$ to be the value measured previously in biochemical studies (25,26), $\tau_1 \approx 1/60\ s$. One possibility is that cooperativity of the receptor-kinase lattice slows down $\tau_1$ to a value closer to what we measure in FRET, $\tau_1 \approx 0.35\ s$. In this case, extrapolating the slow noise to high frequencies does not cause any problems.

To avoid having unphysical noise power at high frequencies, we take the total noise in kinase activity to be a sum of the measured slow noise in Eqn. 82 plus the filtered particle arrival noise in Eqn. 80. There are likely other noise sources at high frequencies, so this modeling choice maximizes our estimate of *E. coli*'s information rate. Since we find that *E. coli* are far from the physical limit, this is a conservative modeling choice. Ultimately, even if we only model noise in kinase activity as being the slow, measurable noise, the effects on the numerical values of the information rate are small.

We will make an additional simplifying assumption. The adaptation time of kinase responses, $\tau_2$, and the correlation time of kinase noise, $\tau_n$, are each roughly $\sim 10\ s$. Therefore, below we will also assume $\tau_2 \approx$

$\tau_n$, which also has small quantitative effects on the results. These simplifications also allow us to derive interpretable analytical expressions.

#### Derivation of the behaviorally-relevant information rate in kinase activity

In this section, we derive the information about current signal encoded in the kinase activity of a typical *E. coli* cell. Here, we seek an expression for the following transfer entropy rate:

$$\dot{I}^{*}_{s\to a} = \lim_{dt\to 0}\frac{1}{dt} I(a(t+dt); s(t)|\{a(t)\})$$
$$= -\left[\partial_\tau I\big(\{a(t)\}; s(t+\tau)\big)\right]_{\tau=0}. \tag{83}$$

Again, the calculation centers on calculating the mutual information between past kinase activity $a$ and signal at some time $\tau$ into the future, $I\big(\{a(t)\}; s(t+\tau)\big)$. The quantity we need to derive this is the posterior distribution of signal given past kinase activity, $P(s(t+\tau)|\{a(t)\})$. Past measurements by us and others (1,24,27) have shown that kinase activity in wild type cells (i.e. cells with all receptor types and with their adaptation system intact) is well-approximated by a Gaussian process. Because of this, and because we consider shallow gradients, we only need the variance of $P(s(t+\tau)|\{a(t)\})$ to compute the mutual information to leading order in $g$ (see the section **Derivation of the behaviorally-relevant information rate in particle arrivals**, above). Thus, we can approximate $s$ and $a$ as jointly Gaussian distributed.

With the approximation that $s$ and $a$ are also jointly Gaussian distributed, $P(s(t+\tau)|\{a(t)\})$ is Gaussian, and therefore we again need to compute a mean $\hat{s}_a(t+\tau)$ and a variance $\sigma^2_{s|a}(\tau)$. Then, the mutual information can then be computed from:

$$I\big(\{a(t)\}; s(t+\tau)\big) = \frac{1}{2}\log\left(\frac{\sigma_s^2}{\sigma^2_{s|a}(\tau)}\right) = -\frac{1}{2}\log\big(1-\rho^2_{as}(\tau)\big), \tag{84}$$

and the predictive information rate is

$$\dot{I}^{*}_{s\to a} = \frac{1}{2}\left[\frac{-\partial_\tau \rho^2_{as}(\tau)}{1-\rho^2_{as}(\tau)}\right]_{\tau=0} \tag{85}$$

Here, $\rho^2_{as}(\tau) = 1 - \frac{\sigma^2_{s|a}(\tau)}{\sigma_s^2}$ is the generalized correlation between $s(t+\tau)$ and past $a$, or the fraction reduction of variance in $s(t+\tau)$ upon observing past $a$.

To compute the rate of information transfer from current signal $s(t)$ to kinase activity $a(t)$, we need the conditional mean and variance of $s(t+\tau)$, $\hat{s}_a(t+\tau)$ and $\sigma^2_{s|a}(\tau)$. These in turn require deriving the kernel $M_a(T)$ that maps past kinase activity $a$ to the conditional mean, $\hat{s}_a(t+\tau)$. This can again be derived using Wiener filtering theory and expressed in terms of the power spectra of $s$ and $a$. These are:

$$S_s(\omega) = F[C_s(T)] = \frac{2\, g^2 \frac{\sigma_v^2}{\tau_v}}{\frac{1}{\tau_v^2} + \omega^2} \tag{86}$$

$$S_a(\omega) = F[C_a(T)] = |K_r(\omega)|^2\, S_r(\omega) + N_{n(\omega)} \tag{87}$$

$$= |K_r(\omega)|^2 \left( r_0^2 \frac{S_s(\omega)}{\omega^2} + r_0 \right) + \frac{2\, D_n}{\frac{1}{\tau_2^2} + \omega^2} \tag{88}$$

$$= \left(\frac{G_r}{\tau_1}\right)^2 \frac{\omega^2}{\left(\frac{1}{\tau_2^2} + \omega^2\right)\left(\left(\frac{1}{\tau_1} + \frac{1}{\tau_2}\right)^2 + \omega^2\right)} \left( r_0^2 \frac{S_s(\omega)}{\omega^2} + r_0 \right) + \frac{2\, D_n}{\frac{1}{\tau_2^2} + \omega^2} \tag{89}$$

$$S_{as}(\omega) = S_{sa}^*(\omega) = F[C_{as}(T)] = -K_r^*(\omega)\, S_{rs}(\omega) \tag{90}$$

$$= -\frac{G_r}{\tau_1} \frac{r_0}{\left(\frac{1}{\tau_2} + i\,\omega\right)\left(\frac{1}{\tau_1} + \frac{1}{\tau_2} + i\,\omega\right)}\, S_s(\omega) \tag{91}$$

where $C_s(T) = \langle s(t)\, s(t+T) \rangle$ , $C_a(T) = \langle (a(t) - a_0)\,(a(t+T) - a_0) \rangle$ , and $C_{as}(T) = \langle (a(t) - a_0)\, s(t+T) \rangle$. The first term in $S_a(\omega)$ comes from responses to signals, the second term comes from filtered particle arrival noise, and the third term comes from internal kinase noise. For convenience, we will define $\tau_3^{-1} = \tau_1^{-1} + \tau_2^{-1}$.

We now need to decompose $S_a(\omega)$ into the product of a causal and an anti-causal part by finding its zeros and poles. The zeros satisfy $S_a(\omega = i\, z_a) = 0$ are complex solutions to the equation:

$$\frac{G_r^2}{\tau_1^2} \left( 2\, r_0^2\, g^2\, \frac{\sigma_v^2}{\tau_v} + r_0\, \omega^2 \left( \frac{1}{\tau_v^2} + \omega^2 \right) \right) + 2\, D_n \left( \frac{1}{\tau_v^2} + \omega^2 \right) \left( \frac{1}{\tau_3^2} + \omega^2 \right) = 0. \tag{92}$$

This can be written in terms of the particle arrival SNR, $\gamma_r = 2\, r_0\, g^2\, \sigma_v^2\, \tau_v^3$, and the ratio of the diffusivity of filtered particle noise and the diffusivity of slow kinase noise, $R = \frac{1}{2} \frac{G_r^2}{\tau_1^2} \frac{r_0}{D_n}$:

$$R\left(\gamma_r + \tau_v^2\, \omega^2\, (1 + \tau_v^2\, \omega^2)\right) + (1 + \tau_v^2\, \omega^2) \left( \frac{\tau_v^2}{\tau_3^2} + \tau_v^2\, \omega^2 \right) = 0 \tag{93}$$

The zeros of $S_a(\omega)$ are:

$$i\, z_{a,1} = i \frac{1}{\tau_v} \frac{1}{\sqrt{2\,(1+R)}} \sqrt{\left(\frac{\tau_v}{\tau_3}\right)^2 + (1+R) - \sqrt{(1+R)\left(1 + R\,(1 - 4\,\gamma_r)\right) - 2\,(1+R)\left(\frac{\tau_v}{\tau_3}\right)^2 + \left(\frac{\tau_v}{\tau_3}\right)^4}},$$

$$i\, z_{a,2} = i \frac{1}{\tau_v} \frac{1}{\sqrt{2\,(1+R)}} \sqrt{\left(\frac{\tau_v}{\tau_3}\right)^2 + (1+R) + \sqrt{(1+R)\left(1 + R\,(1 - 4\,\gamma_r)\right) - 2\,(1+R)\left(\frac{\tau_v}{\tau_3}\right)^2 + \left(\frac{\tau_v}{\tau_3}\right)^4}}, \tag{94}$$

as well as their complex conjugates.

The poles of $S_a(\omega)$ satisfy $\frac{1}{S_a(\omega=i\,p_a)}=0$ and are $i\,p_{a,1}=i\frac{1}{\tau_v}$, $i\,p_{a,2}=i\frac{1}{\tau_2}$, and , $i\,p_{a,3}=i\frac{1}{\tau_3}$, as well as their complex conjugates.

We decompose $S_a(\omega)$ as:

$$S_a(\omega)=\phi_a(\omega)\,\phi_a^*(\omega) \tag{95}$$

where

$$\phi_a(\omega)=\sqrt{2\,D_n\,(1+R)}\frac{(z_{a,1}-i\,\omega)(z_{a,2}-i\,\omega)}{(p_{a,1}-i\,\omega)\,(p_{a,2}-i\,\omega)\,(p_{a,3}-i\,\omega)}. \tag{96}$$

Next, we need the causal part of the following (see Appendix A):

$$\frac{S_{as}(\omega)}{\phi_a^*(\omega)}e^{-i\,\omega\,\tau}=-\frac{G_r/\tau_1}{\sqrt{2\,D_n\,(1+R)}}\frac{2\,r_0\,g^2\,\sigma_v^2\frac{1}{\tau_v}}{\left(\frac{1}{\tau_v}-i\,\omega\right)\,(z_{a,1}+i\,\omega)\,(z_{a,2}+i\,\omega)}e^{-i\,\omega\,\tau} \tag{97}$$

Again, we find the causal part of this expression by doing a partial fraction decomposition and keeping only the terms with poles and zeros that have negative imaginary part:

$$\frac{S_{as}(\omega)}{\phi_a^*(\omega)}e^{-i\,\omega\,\tau}=\frac{A}{\left(\frac{1}{\tau_v}-i\,\omega\right)}+\frac{B}{(z_{a,1}+i\,\omega)}+\frac{C}{(z_{a,2}+i\,\omega)}, \tag{98}$$

for unknown $A$, $B$, and $C$. Only the pole of the first term (at $\omega=-i\frac{1}{\tau_v}$) has negative imaginary part, so we only need to compute $A$ to get the causal part of this expression. This is:

$$A=\left[-\frac{G_r/\tau_1}{\sqrt{2\,D_n\,(1+R)}}\frac{2\,r_0\,g^2\,\sigma_v^2\frac{1}{\tau_v}}{(z_{a,1}+i\,\omega)\,(z_{a,2}+i\,\omega)}e^{-i\,\omega\,\tau}\right]_{\omega=-i\frac{1}{\tau_v}} \tag{99}$$

$$=-\frac{G_r/\tau_1}{\sqrt{2\,D_n\,(1+R)}}\frac{2\,r_0\,g^2\,\sigma_v^2\,\tau_v}{(1+z_{a,1}\,\tau_v)(1+z_{a,2}\,\tau_v)}e^{-\frac{\tau}{\tau_v}}, \tag{100}$$

and the causal part of $S_{as}(\omega)/\phi_a^*(\omega)$ is then:

$$\left[\frac{S_{as}(\omega)}{\phi_a^*(\omega)}e^{-i\,\omega\,\tau}\right]^+=-\frac{G_r/\tau_1}{\sqrt{2\,D_n\,(1+R)}}\frac{2\,r_0\,g^2\,\sigma_v^2\,\tau_v}{(1+z_{a,1}\,\tau_v)(1+z_{a,2}\,\tau_v)}\frac{e^{-\frac{\tau}{\tau_v}}}{\left(\frac{1}{\tau_v}-i\,\omega\right)}. \tag{101}$$

Finally, like $C_{rs}(\tau)$ in the section above, $C_{as}(\tau)\propto\exp\left(-\frac{\tau}{\tau_v}\right)$ when $\tau\geq 0$.

With these expressions, the optimal kernel that computes the mean of $p(s(t+\tau)|\{a\})$ is (Appendix A):

$$M_a(\omega)=\frac{1}{\phi_a(\omega)}\left[\frac{S_{as}(\omega)}{\phi_a^*(\omega)}e^{-i\,\omega\,\tau}\right]^+ \tag{102}$$

$$= -e^{-\frac{\tau}{\tau_v}} \frac{2\frac{G_r}{\tau_1} r_0 \, g^2 \, \sigma_v^2 \, \tau_v}{2\, D_n \, (1+R) \left(1 + z_{a,1}\,\tau_v\right)\left(1 + z_{a,2}\,\tau_v\right)} \frac{\left(\frac{1}{\tau_2} - i\,\omega\right)\left(\frac{1}{\tau_3} - i\,\omega\right)}{\left(z_{a,1} - i\,\omega\right)\left(z_{a,2} - i\,\omega\right)} \quad (103)$$

We discuss this kernel in the following section.

The variance of $P(s(t+\tau)|\{a\})$, $\sigma_{s|a}^2(\tau)$, is (Appendix A, Eqn. 181):

$$\sigma_{s|a}^2(\tau) = \sigma_s^2 - \frac{1}{2\pi}\int_{-\infty}^{\infty} S_{as}^*(\omega)\, e^{i\,\omega\,\tau}\, M_a(\omega)\, d\omega \quad (104)$$

$$= \sigma_s^2 \left( 1 - e^{-2\frac{\tau}{\tau_v}} \frac{2\frac{G_r^2}{\tau_1^2} r_0^2 \, g^2 \, \sigma_v^2 \, \tau_v^3}{2\, D_n \, (1+R)\left(1 + z_{a,1}\,\tau_v\right)^2 \left(1 + z_{a,2}\,\tau_v\right)^2} \right), \quad (105)$$

where $\sigma_s^2 = g^2 \, \sigma_v^2$. Therefore, the correlation coefficient $\rho_{as}^2(\tau)$ is:

$$\rho_{as}^2(\tau) = 1 - \frac{\sigma_{s|a}^2(\tau)}{\sigma_s^2} = e^{-2\frac{\tau}{\tau_v}} \frac{2\frac{G_r^2}{\tau_1^2} r_0^2 \, g^2 \, \sigma_v^2 \, \tau_v^3}{2\, D_n \, (1+R)\left(1 + z_{a,1}\,\tau_v\right)^2 \left(1 + z_{a,2}\,\tau_v\right)^2}, \quad (106)$$

or in terms of $\gamma_r = 2\, r_0 \, g^2 \, \sigma_v^2 \, \tau_v^3$ and $R = \frac{1}{2}\frac{G_r^2}{\tau_1^2}\frac{r_0}{D_n}$:

$$= e^{-2\frac{\tau}{\tau_v}} \frac{R}{(1+R)} \frac{\gamma_r}{\left(1 + z_{a,1}\,\tau_v\right)^2 \left(1 + z_{a,2}\,\tau_v\right)^2}. \quad (107)$$

Finally, using Eqn. 85 above, we find that the information about current signal encoded in *E. coli*'s kinase activity is:

$$\dot{I}_{s\to a}^* = \frac{1}{\tau_v}\frac{\rho_{as}^2(\tau=0)}{1-\rho_{as}^2(\tau=0)} = \frac{1}{\tau_v} \frac{\frac{R}{(1+R)}\frac{\gamma_r}{\left(1 + z_{a,1}\,\tau_v\right)^2 \left(1 + z_{a,2}\,\tau_v\right)^2}}{1 - \frac{R}{(1+R)}\frac{\gamma_r}{\left(1 + z_{a,1}\,\tau_v\right)^2 \left(1 + z_{a,2}\,\tau_v\right)^2}}. \quad (108)$$

In shallow gradients, $\dot{I}_{s\to a}^* \approx \frac{1}{\tau_v}\rho_{as}^2(\tau=0) \approx \frac{2}{\tau_v} I\left(\{a(t)\}; s(t)\right)$, and only the leading order $g^2$ term of $\dot{I}_{s\to a}^*$ contributes to the final expression. Since $\gamma_r \propto g^2$ in Eqn. 107, we can get the shallow-gradient expression for $\dot{I}_{s\to a}^*$ by evaluating $z_{a,1}$ and $z_{a,2}$ at $g = 0$. This is equivalent to taking $\gamma_r \to 0$, which gives:

$$z_{a,1} \approx \frac{1}{\tau_3\,\sqrt{1+R}}, \qquad z_{a,2} \approx \frac{1}{\tau_v}. \quad (109)$$

Thus, in shallow gradients, we get:

$$\dot{I}_{s\to a}^* \approx \frac{1}{\tau_v}\frac{1}{4}\frac{R}{1+R}\frac{\gamma_r}{\left(1 + \frac{\tau_v}{\tau_3}\frac{1}{\sqrt{1+R}}\right)^2} = \frac{1}{\tau_v}\frac{1}{4}\frac{\frac{1}{2}\frac{G_r^2}{\tau_1^2}\frac{r_0}{D_n}}{1+\frac{1}{2}\frac{G_r^2}{\tau_1^2}\frac{r_0}{D_n}} \frac{2\, r_0 \, g^2 \, \sigma_v^2 \, \tau_v^3}{\left(1 + \frac{\tau_v}{\tau_3}\left(1 + \frac{1}{2}\frac{G_r^2}{\tau_1^2}\frac{r_0}{D_n}\right)^{-1/2}\right)^2}, \quad (110)$$

where again $\tau_3^{-1} = \tau_1^{-1} + \tau_2^{-1}$. Furthermore, since $\tau_1 \ll \tau_v$, taking $\tau_1 \to 0$ only slightly increases the information rate and gives a simpler expression in terms of a kinase signal to noise ratio, $\gamma_a = \frac{G_r^2}{D_n} r_0^2\, g^2\, \sigma_v^2\, \tau_v$, and the particle arrival signal to noise ratio, $\gamma_r = 2\, r_0\, g^2 \sigma_v^2\, \tau_v^3$:

$$\dot{I}_{s\to a}^{*} \approx \frac{1}{\tau_v}\frac{1}{4}\,\gamma_a \frac{\frac{\gamma_r}{\gamma_a}}{\left(1+\sqrt{\frac{\gamma_r}{\gamma_a}}\right)^2} = \frac{1}{\tau_v}\frac{1}{4}\frac{G_r^2}{D_n} r_0^2\, g^2\, \sigma_v^2\, \tau_v \frac{\frac{2\, D_n\, \tau_v^2}{G_r^2\, r_0}}{\left(1+\sqrt{\frac{2\, D_n\, \tau_v^2}{G_r^2\, r_0}}\right)^2}. \tag{111}$$

We also note that for finite $g$ but $\tau_1 \to 0$, $z_{a,1}$ and $z_{a,2}$ are:

$$z_{a,1} \approx \frac{1}{\tau_v}\frac{1}{\sqrt{2}}\sqrt{1+\frac{\gamma_r}{\gamma_a} - \sqrt{1 - 4\,\gamma_r - 2\frac{\gamma_r}{\gamma_a} + \left(\frac{\gamma_r}{\gamma_a}\right)^2}},$$

$$z_{a,2} \approx \frac{1}{\tau_v}\frac{1}{\sqrt{2}}\sqrt{1+\frac{\gamma_r}{\gamma_a} + \sqrt{1 - 4\,\gamma_r - 2\frac{\gamma_r}{\gamma_a} + \left(\frac{\gamma_r}{\gamma_a}\right)^2}}. \tag{112}$$

We plugged these expressions into Eqn. 108, with $\frac{R}{1+R} \to 1$ as $\tau_1 \to 0$, to generate the plots in Fig. 3 of the main text.

Eqns. 46, 54, and 108 for the information rates $\dot{I}_{s\to r}^{*}$ and $\dot{I}_{s\to a}^{*}$ are nearly exact, but make several assumptions. They require $r_0\, \tau_v \gg 1$ so that we can approximate particle arrivals as Gaussian. They also use Gaussian approximations for the mutual information quantities $I(s(t);\{r\})$ and $I(s(t);\{a\})$, which are valid when these quantities are small (shallow gradients, small $g$). We used linear theory to model kinase responses, which is valid if deviations in kinase activity from baseline are small—i.e. when $g$ is small. And we ignored feedbacks in which responses to signals change the signal statistics that the cell experiences, again valid when $g$ is small. Each of these assumptions can break at a different characteristic value of $g$: for particle arrival rate, small $g$ means $\gamma_r \ll 1$; for kinase activity, small $g$ means $\gamma_a \ll 1$. That all said, Eqns. 46, 54, and 108 currently provide our best analytical insight into information transfer during chemotaxis.

#### Optimal kernel for estimating signal from kinase activity

To understand the kernel $M_a(\omega)$ that constructs an estimate of the current signal, $s(t)$, from past kinase activity, $\{a\}$, we first multiply it by the kinase response function of particle arrivals, $K_r(\omega)$. This gives a composite kernel that effectively maps the past of particle arrivals $r$, corrupted by kinase noise, to an estimate of the signal $s(t)$:

$$-M_a(\omega)\, K_r(\omega) = -\frac{2\left(\frac{G_r}{\tau_1}\right)^2 r_0\, g^2\, \sigma_v^2\, \tau_v}{2\, D_n\,(1+R)\,\big(1+z_{a,1}\,\tau_v\big)\big(1+z_{a,2}\,\tau_v\big)} \frac{(-i\,\omega)}{\big(z_{a,1} - i\,\omega\big)\big(z_{a,2} - i\,\omega\big)}. \tag{113}$$

In the time domain, this is:

$$IFT[-M_a(\omega)\, K_r(\omega)] = \frac{2\left(\frac{G_r}{\tau_1}\right)^2 r_0\, g^2\, \sigma_v^2\, \tau_v}{2\, D_n\, (1+R)} \frac{\left(z_{a,2}\, \exp(-z_{a,2}\, t) - z_{a,1}\, \exp(-z_{a,1}\, t)\right)}{(1+z_{a,1}\,\tau_v)(1+z_{a,2}\,\tau_v)(z_{a,1}-z_{a,2})}\, \Theta(t), \quad (114)$$

This composite kernel that effectively acts on particle arrivals has the same structure as the optimal kernel $M_r(T)$ (Eqn. 55) for directly constructing $s(t)$ from particle arrivals. It's biphasic and adapts perfectly, although with different time scales than $M_r(T)$. This means that $M_a(T)$ attempts to invert the kinase response function $K_r(T)$, to the extent possible given the kinase noise $N_n(T)$, and then apply something as close as possible to the optimal kernel for particle counts, $M_r(T)$.

Taking this further, we argue that the optimal kernel acting on particle counts, $M_r(T)$, is the response kernel that the cell should *try* to implement as its behavioral output using its intracellular signaling network (up to changes of units). However, the cell has to communicate information about the signal $s(t)$ through multiple chemical species in order to send them from the kinases at one location to the motors at various other locations. These steps impose constraints on the cell's signaling pathway, and they add noise. Despite this, the cell should be attempting to make its composite kernel from input (particle counts) to output (tumble rate) look like $M_r(T)$.

#### Simulation details

In Figs. 4A and 4B of the main text, we performed two types of simulations in which a run-tumble particle moved in three dimensions with rotational diffusion through a concentration gradient. In both cases, we compare "ideal" cells, which sense particle arrivals directly, to "*E. coli*," which respond to particle arrivals with changes in kinase activity. In Fig. 4A, we compute the optimal estimates of the signal in each case, $\hat{s}_r(t)$ and $\hat{s}_a(t)$, but the cells' tumble behaviors do not depend on the signal. Optimal estimates of $s(t)$ were computed using the kernels above. In Fig. 4B, the cells' tumble rates are modulated by their optimal estimates of the signal. We detail these two simulations below.

In both Fig. 4A and 4B, the velocity correlation time $\tau_v$ has contributions from the cell's average tumble rate, $\lambda_0$, the persistence of tumbles, $\alpha$, and rotational diffusion, $D_r$: $\tau_v^{-1} = (1-\alpha)\,\lambda_{R0} + 2\, D_r$. We took $\alpha = 0$ and $D_r = 0.044\ \mathrm{rad}^2/\mathrm{s}$ based on our previous measurements (1), and then computed the tumble rate $\lambda_0$ from the value of $\tau_v$ measured here. Assuming nearly instantaneous tumbles, we computed the swimming speed to use in simulations from $\sigma_v^2 = v_0^2/3$ in 3D.

Tumbles were generated by sampling a uniform random variable $u$, and a tumble occurred if $u < \lambda_0\, \Delta t$. Tumbles lasted one time step. Rotational diffusion during runs was simulated as before (28,29). The simulation was performed by stepping forward in time by discrete steps of size $\Delta t = \min(\tau_v,\ k_1^{-1})/100$.

Cells were initialized at position $x(t=0) = 0$ in a concentration gradient $c(x) = c_0 \exp(g\, x)$. At each time step, stochastic molecule arrival rate was computed from $r(t) = k_D\, c(x(t)) + \sqrt{r_0/dt}\, \eta(t)$, where $\eta(t)$ was a standard Gaussian-distributed random variable simulating shot noise. Kinase activity was computed after the simulation using Eqn. 4 of the main text, with $\tau_1 \to 0$ and otherwise measured parameters (Fig. S1), and the kinase noise $\eta_n(t)$ was simulated separately. Note that any nonlinear model of kinase activity must behave like the measured linear response kernel and noise when the signals are small.

**Simulations without tumble responses to signals (Fig. 4A)**

In Fig. 4A (no tumble response to signals), we simulated run-and-tumble motion with rotational diffusion in a static gradient, as described above, to get $s(t)$, $c(t)$, and $r(t)$. After the simulation, we computed $a(t)$ and the optimal estimates of the signal $s(t)$, $\hat{s}_r(t) = \int_{-\infty}^{t} M_r(t-t')\,(r(t')-r_0)\,dt'$ and $\hat{s}_a(t) = \int_{-\infty}^{t} M_a(t-t')\,(a(t')-a_0)\,dt'$, using expressions above. We used these simulations to check our expressions for the correlation coefficients between the optimal signal estimates and the true signals, $\rho_{rs}^2$ and $\rho_{as}^2$, and thus validate our expressions for the information rates. Specifically, we numerically computed these correlations in the simulations as $\rho_{rs}^2 = \langle s(t)\,\hat{s}_r(t)\rangle^2 / \left(\sigma_s^2\,\sigma_{\hat{s}_r}^2\right)$ and $\rho_{as}^2 = \langle s(t)\,\hat{s}_a(t)\rangle^2 / \left(\sigma_s^2\,\sigma_{\hat{s}_a}^2\right)$ and compared to the theoretical expressions. For this, we simulated 5000 cells for $T = 100\,\tau_v$ and excluded the initial transient ($10\,\tau_v$) from the analysis. Fig. S5 below shows excellent agreement between the theory and simulations.

**Simulations with tumble responses to signals (Fig. 4BC)**

Our main goals were to construct a model that allowed fair comparison between "ideal" and "realistic" cells, and to demonstrate that the loss of information due to kinase noise translates into a reduction of drift speed.

To simulate tumbling in response to estimated signals in Fig. 4B, we needed to simulate kinase activity $a(t)$ and the optimal estimates of the signal. To achieve this, we next derive dynamical systems for $a(t)$, $\hat{s}_r(t)$, and $\hat{s}_a(t)$. In general, if a quantity is the convolution of some variable, $h(t)$, with a kernel $g(T)$:

$$f(t) = \int_{-\infty}^{t} g(t-t')\,h(t')\,dt', \tag{115}$$

then its time derivative is:

$$\frac{df}{dt} = g(0)\,h(t) + \int_{-\infty}^{t} g'(t-t')\,h(t')\,dt'. \tag{116}$$

First consider kinase activity,

$$a(t) = a_0 - \int_{-\infty}^{t} K_a(t-t')\,(r(t')-r_0)\,dt' + \eta_n(t). \tag{117}$$

As $\tau_1 \to 0$, this becomes

$$a(t) = a_0 - G_r\left((r(t)-r_0) - \frac{1}{\tau_2}\int_{-\infty}^{t} \exp\left(-\frac{(t-t')}{\tau_2}\right)(r(t')-r_0)\,dt'\right) + \eta_n(t). \tag{118}$$

We can write down a dynamical system for the deterministic part of $a(t)$ by defining the convolution term above as a new variable:

$$a_r(t) = \frac{G_r}{\tau_2}\int_{-\infty}^{t} \exp\left(-\frac{(t-t')}{\tau_2}\right)(r(t')-r_0)\,dt'. \tag{119}$$

Then, using Eqn. 116, its time derivative is:

$$\frac{d}{dt}a_r(t) = \frac{G_r}{\tau_2}\,(r(t) - r_0) - \frac{1}{\tau_2}a_r(t). \tag{120}$$

The internal noise $\eta_n(t)$ is an Ornstein-Uhlenbeck process with diffusivity $D_n$ and time scale $\tau_n \approx \tau_2$, which we simulated using standard methods. Then $a(t)$ at each time step was computed from the dynamical variables $r(t)$, $a_r(t)$, and $\eta(t)$ using:

$$a(t) = a_0 - G_r(r(t) - r_0) + a_r(t) + \eta_n(t). \tag{121}$$

To simulate $\hat{s}_r(t)$, we first use its definition, $\hat{s}_r(t) = \int_{-\infty}^{t} M_r(t - t')\,(r(t') - r_0)\,dt'$, where the optimal kernel $M_r(T)$ is (Eqn. 59, $\gamma > 1/4$):

$$M_r(t) = A\exp(-k_1\,t)\,\left(\cos(k_2\,t) - \frac{k_1}{k_2}\sin(k_2\,t)\right),$$

and

$$A = \frac{\gamma_r}{r_0\,\tau_v^2}\frac{1}{\left(1 + \sqrt{1 + 2\sqrt{\gamma_r}} + \sqrt{\gamma_r}\right)}, \tag{122}$$

$$k_1 = \frac{1}{\tau_v}\frac{1}{2}\sqrt{2\sqrt{\gamma_r} + 1}, \tag{123}$$

$$k_2 = \frac{1}{\tau_v}\frac{1}{2}\sqrt{2\sqrt{\gamma_r} - 1}. \tag{124}$$

We can write down a dynamical system to simulate $\hat{s}_r(t)$ by defining the following two variables:

$$\hat{s}_r^1(t) = A\int_{-\infty}^{t} \exp\big(-k_1\,(t - t')\big)\cos\big(k_2\,(t - t')\big)\,\,(r(t') - r_0)\,dt', \tag{125}$$

$$\hat{s}_r^2(t) = A\int_{-\infty}^{t} \exp\big(-k_1\,(t - t')\big)\sin\big(k_2\,(t - t')\big)\,\,(r(t') - r_0)\,dt', \tag{126}$$

and using $\hat{s}_r(t) = \hat{s}_r^1(t) - \frac{k_1}{k_2}\hat{s}_r^2(t)$.

We can derive a closed system of equations for the dynamics of these variables using Eqn. 116:

$$\frac{d}{dt}\hat{s}_r^1(t) = A\,(r(t) - r_0) - k_1\,\hat{s}_r^1(t) - k_2\,\hat{s}_r^2(t), \tag{127}$$

$$\frac{d}{dt}\hat{s}_r^2(t) = k_2\,\hat{s}_r^1(t) - k_1\,\hat{s}_r^2(t). \tag{128}$$

A similar approach can be used to derive dynamics $\hat{s}_a(t) = \int_{-\infty}^{t} M_a(t - t')\,(a(t') - a_0)\,dt'$. For $\tau_1 \ll \tau_v$ and $\tau_n \approx \tau_2$, which are consistent with our measurements, $M_a(t)$ has the form (inverse Fourier transform of Eqn. 103, after taking parameter limits):

$$M_a(t) = -(B_3\exp(-k_3\,t) - B_4\exp(-k_4\,t)), \tag{129}$$

where $k_3 = z_{a,1}$, $k_4 = z_{a,2}$ (Eqn. 94), and

$$B_3 = \frac{2\,g^2\,\sigma_v^2\,\tau_v}{G_r}\frac{\left(z_{a,1} - \frac{1}{\tau_2}\right)}{\left(z_{a,1} - z_{a,2}\right)\left(1 + z_{a,1}\,\tau_v\right)\left(1 + z_{a,2}\,\tau_v\right)}, \tag{130}$$

$$B_4 = \frac{2\,g^2\,\sigma_v^2\,\tau_v}{G_r}\frac{\left(z_{a,2} - \frac{1}{\tau_2}\right)}{\left(z_{a,1} - z_{a,2}\right)\left(1 + z_{a,1}\,\tau_v\right)\left(1 + z_{a,2}\,\tau_v\right)}. \tag{131}$$

Next, we define the following variables:

$$\hat{s}_a^1(t) = -B_3 \int_{-\infty}^{t} \exp\left(-k_3\,(t - t')\right)\left(a(t') - a_0\right)dt', \tag{132}$$

$$\hat{s}_a^2(t) = -B_4 \int_{-\infty}^{t} \exp\left(-k_4\,(t - t')\right)\left(a(t') - a_0\right)dt', \tag{133}$$

and use $\hat{s}_a(t) = \hat{s}_a^1(t) - \hat{s}_a^2(t)$.

By similar steps as above, the dynamics of these variables are:

$$\frac{d}{dt}\hat{s}_a^1(t) = -B_3\,\left(a(t') - a_0\right) - k_3\,\hat{s}_a^1(t), \tag{134}$$

$$\frac{d}{dt}s_a^2(t) = -B_4\,\left(a(t') - a_0\right) - k_4\,\hat{s}_a^2(t). \tag{135}$$

Next, the optimal estimates of signals, $\hat{s}_r(t)$ and $\hat{s}_a(t)$, modulate the cells' tumble rates. For both ideal and realistic cells, we again modeled tumbling as a Poisson process, but with time-varying rate depending on the estimated signal from their available observations, $\lambda(t) = \lambda\left(\hat{s}_r(t)\right)$ or $\lambda(t) = \lambda\left(\hat{s}_a(t)\right)$. Tumbling is a stochastic process, and the "signal to noise" ratio of this process depends on the variance of the input that drives the tumble rate. To allow fair comparisons between the ideal and realistic cells, we modeled the tumble rates as functions of rescaled optimal estimators $\hat{s}_r(t)$ and $\hat{s}_a(t)$ such that the two estimators had the same variance, which was also independent of $g$:

$$\lambda(t) = \lambda\left(\frac{\hat{s}(t)}{v_0\,g\,\rho}\right), \tag{136}$$

where $\hat{s}(t) = \hat{s}_r(t)$ and $\rho = \rho_{rs}$ for ideal cells, and $\hat{s}(t) = \hat{s}_a(t)$ and $\rho = \rho_{as}$ for realistic cells. Rescaling the optimal estimates of the signal does not change the information rates. The factor of $\rho$ corrects for the fact the posterior mean, $\hat{s}(t)$, is shrunken towards zero by a factor of $\rho$. Note that our goal here is not to develop a mechanistic biochemical model, which has been pursued elsewhere (28,30–35). Rather, we aim to examine the effects of information on gradient climbing in a minimal setting that allows fair comparison between ideal cells, with access to information $\dot{I}_{s\to r}^*$, and *E. coli*-like cells, with access to information $\dot{I}_{s\to a}^*$.

Defining $y(t) = \frac{\hat{s}(t)}{v_0\,g\,\rho}$ for short, the tumble rate was parameterized as follows:

$$\lambda(t) = \lambda_0 \frac{A(G_m, \Delta)}{1 + \exp(B(G_m, \Delta)\,G_m\,y(t) + \Delta)}. \tag{137}$$

The constants $A(G_m, \Delta)$ and $B(G_m, \Delta)$ were determined by enforcing the baseline tumble rate in the absence of signal $\lambda(y = 0) = \lambda_0$, and by enforcing that the linearized tumble rate for small signals was always

$$\lambda(t) \approx \lambda_0 \left(1 - G_m\, y(t)\right). \tag{138}$$

Thus, $G_m$ is the gain of the tumble response to the input $y(t)$. The offset parameter $\Delta$ determined where on the "tumble curve" cells lie at baseline, and also determined the maximum possible tumble rate when $y \ll 0$. For example, $\Delta \to \infty$ gives $\lambda(t) = \lambda_0 \exp\left(-G_m\, y(t)\right)$, a functional form that has been used before by others (e.g. (30)). In general,

$$A = 1 + \exp(\Delta) \tag{139}$$

$$B = \frac{1 + \exp(\Delta)}{\exp(\Delta)} \tag{140}$$

We chose $\Delta = 1$ and $G_m = 4$ so that the chemotaxis coefficient, $\chi$, of the "realistic" cells was comparable to values we measured previously for *E. coli* climbing gradients of methyl-aspartate (1).

We simulated $N_{cells} = 10{,}000$ cells for $T = 100\,\tau_v$ and excluded the initial transient ($10\,\tau_v$) from the analysis. For the smallest value of $g = 0.05\ \mathrm{mm}^{-1}$, we simulated $N_{cells} = 50{,}000$ cells. Drift speeds were computed as the average up-gradient velocity over time and cells, $v_d = \langle v_x(t) \rangle$, and standard errors were computed as $\sigma_{v_d}^2 = Var\left(v_x(t)\right)/(N_{cells}\, T/\tau_v)$, where the denominator is approximately the number of independent samples. Information rates were computed from simulations by computing the correlation coefficient between the true and estimated signals and plugging into Eqn. 46 or Eqn. 108, e.g. $\dot{I} = \frac{1}{\tau_v} \frac{\rho^2}{1-\rho^2}$. The numerically-computed information rates agreed with our theoretical expressions in shallow gradients, where feedback from signals onto behavior was weak (29). In steeper simulated gradients, the information rate in simulations was larger than in our theory, up to a factor of 2 for ideal cells in $g = 0.4\ \mathrm{mm}^{-1}$, mostly because the actual $\tau_v$ in simulation dynamically became longer than the value we input to the simulation.

#### Estimating population variability in $\boldsymbol{\eta}$

Although the median phenotype in our strain's population is far from the molecule-counting limit on chemical sensing, individual cells with the same genes exhibit large non-genetic differences. Therefore, it's possible that a subset of the population could approach the bound. To tentatively test this possibility, we use maximum likelihood estimation.

Due to variations in swimming, kinase response, and kinase noise parameters among cells, there is a distribution of $\eta = \dot{I}_{s\to a}^{*} / \dot{I}_{s\to r}^{*}$ in the population when exposed to background concentration $c_0$ and gradient steepness $g$. If we could measure all parameters in the same individual cells, we could try to construct this distribution directly. Instead, our experimental setup gives single-cell swimming, response, and noise parameters in different cells. With these single-cell parameters, we can make tentative inferences about the variation in $\eta$ in the population if we assume that swimming, response, and noise parameters are uncorrelated in single cells. This is not necessarily true—for example, cells with higher kinase gain, $G_r$, might also have larger kinase fluctuations, $\sigma_n^2$—but it gives a first estimate. Our single-cell parameters also have uncertainties, and we want to account for this in our estimation.

Using the data we have, we can draw $N$ sample parameters and their uncertainties from single cells in our data set. We chose $N \approx 100$ in a given $c_0$ condition to be the smaller of the number of cells measured in our kinase response experiment in that condition and the number of cells measured in our kinase noise experiment in that condition. In particular, we draw all kinase response parameters are taken from one measured cell and all noise parameters from another measured cell. For swimming parameters, we sample a value of tumble bias from the TB distribution and take the average run duration and swimming speed associated with that tumble bias.

For each sample parameter set $i$, we compute a noise-corrupted estimate of $\eta_i$, where noise comes from the uncertainties in the single-cell parameters. When we transform the noisy observations $\eta_i \to y_i = \log\left(\frac{\eta_i}{1-\eta_i}\right)$, empirically the collection of sampled values of $y_i \in (-\infty, \infty)$ is roughly Gaussian distributed. Therefore, we assume a model in which $y_i = x_i + \xi_i$, where $x_i$ is the true single-cell value, which is Gaussian-distributed due to phenotypic variation in the population with mean $\mu$ and variance $S$, and $\xi_i$ is zero-mean Gaussian noise due to our parameter uncertainties with variance $\sigma_i^2$. Since we know the parameter uncertainties, we know $\sigma_i^2$ for each sample $y_i$, from which we seek to infer $\mu$ and $S$.

Mathematically, we have:

$$P(\{y\},\{x\}|\mu,S) = \prod_{i=1}^{N} P(y_i|x_i)\, P(x_i|\mu,S) \tag{141}$$

$$= \prod_{i=1}^{N} \frac{1}{\sqrt{2\pi\sigma_i^2}} \exp\left(-\frac{1}{2}\frac{(y_i - x_i)^2}{\sigma_i^2}\right) \frac{1}{\sqrt{2\pi S}} \exp\left(-\frac{1}{2}\frac{(x_i - \mu)^2}{S}\right). \tag{142}$$

$P(x_i|\mu,S)$ is the probability density of a single cell from the population having phenotype $x_i$. $P(y_i|x_i)$ is the probability distribution of the noise added due to our parameter uncertainties, resulting in observation $y_i$.

Since we only observe $\{y\}$, and not the true phenotypes $\{x\}$, we marginalize out $\{x\}$:

$$P(\{y\}|\mu,S) = \int P(\{y\},\{x\}|\mu,S)\, d\{x\} = \prod_{i=1}^{N} P(y_i|\mu,S) \tag{143}$$

$$= \prod_{i=1}^{N} \frac{1}{\sqrt{2\pi\left(\sigma_i^2 + S\right)}} \exp\left(-\frac{1}{2}\frac{(y_i - \mu)^2}{\sigma_i^2 + S}\right). \tag{144}$$

This is the likelihood of seeing noise-corrupted phenotypes $\{y\}$ if the true parameters were $\mu$ and $S$. With this, we can estimate the population mean $\mu$ and variance $S$ by maximum likelihood estimation. The log-likelihood of the data $\{y\}$ is:

$$L(\mu, S) = \log\big(P(\{y\}|\mu, S)\big) = -\frac{1}{2}\sum_{i=1}^{N}\left(\log\left(2\pi(\sigma_i^2 + S)\right) + \frac{(y_i - \mu)^2}{\sigma_i^2 + S}\right). \quad (145)$$

To find the parameter values that maximize the log likelihood, we first take derivative with respect to $\mu$ set it to zero:

$$\frac{d}{d\mu}L(\mu, S) = \sum_{i=1}^{N}\frac{(y_i - \mu)}{\sigma_i^2 + S} = 0. \quad (146)$$

Solving for $\mu$:

$$\mu = \frac{\sum_{i=1}^{N}\frac{y_i}{\sigma_i^2 + S}}{\sum_{i=1}^{N}\frac{1}{\sigma_i^2 + S}}. \quad (147)$$

The derivative of the log likelihood with respect to $S$ gives a second equation:

$$\frac{d}{dS}L(\mu, S) = -\frac{1}{2}\sum_{i=1}^{N}\left(\frac{1}{\sigma_i^2 + S} - \frac{(y_i - \mu)^2}{\left(\sigma_i^2 + S\right)^2}\right) = 0. \quad (148)$$

This equation cannot be solved for $S$ analytically, but we can numerically solve it together with Eqn. 147 to get estimates of $\mu$ and $S$. Table S1 shows estimates of $\mu$ and $S$ in each $(c_0, g)$ condition. Point estimates are the mean estimated parameters from 100 bootstrapped data sets of single-cell phenotypes $\{y\}$, and error bars are the standard deviations.

| $g \rightarrow$ <br> $c_0 \downarrow$ | $0^+$ mm$^{-1}$ | 0.1 mm$^{-1}$ | 0.2 mm$^{-1}$ | 0.3 mm$^{-1}$ | 0.4 mm$^{-1}$ |
|---|---|---|---|---|---|
| 0.1 μM | $\mu = -5.8 \pm 0.1$ <br> $S = 1.6 \pm 0.2$ | $\mu = -5.4 \pm 0.1$ <br> $S = 1.7 \pm 0.2$ | $\mu = -5.3 \pm 0.1$ <br> $S = 1.7 \pm 0.2$ | $\mu = -4.9 \pm 0.2$ <br> $S = 1.7 \pm 0.3$ | $\mu = -4.7 \pm 0.1$ <br> $S = 1.6 \pm 0.3$ |
| 1 μM | $\mu = -4.4 \pm 0.1$ <br> $S = 1.0 \pm 0.1$ | $\mu = -3.4 \pm 0.1$ <br> $S = 1.0 \pm 0.2$ | $\mu = -2.8 \pm 0.1$ <br> $S = 1.0 \pm 0.2$ | $\mu = -2.4 \pm 0.1$ <br> $S = 1.1 \pm 0.2$ | $\mu = -2.1 \pm 0.1$ <br> $S = 1.0 \pm 0.2$ |
| 10 μM | $\mu = -6.1 \pm 0.1$ <br> $S = 0.8 \pm 0.1$ | $\mu = -4.1 \pm 0.1$ <br> $S = 0.8 \pm 0.2$ | $\mu = -3.3 \pm 0.1$ <br> $S = 0.8 \pm 0.2$ | $\mu = -2.8 \pm 0.1$ <br> $S = 0.8 \pm 0.1$ | $\mu = -2.5 \pm 0.1$ <br> $S = 0.9 \pm 0.2$ |

**Table S1: Parameters characterizing population variability in $\eta$ from maximum-likelihood estimation.**

To estimate population variation in $\eta$, we transform the Gaussian distribution of $x = \log\left(\frac{\eta}{1-\eta}\right)$ to the distribution of $\eta = \frac{\exp(x)}{1+\exp(x)}$ (abusing notation in the sense that this $\eta$ is noise-free):

$$P(\eta) = P\big(x(\eta)\big)\left|\frac{dx(\eta)}{d\eta}\right| = \frac{1}{\eta\,(1-\eta)}\frac{1}{\sqrt{2\pi S}}\exp\left(-\frac{1}{2}\frac{\left(\log\left(\frac{\eta}{1-\eta}\right) - \mu\right)^2}{S}\right). \quad (149)$$

The larger error bars in Fig. 3C of the main text were computed from the 5% and 95% percentiles of this distribution in each $(c_0, g)$ condition.

#### Information about current versus past signals encoded in kinase activity

We previously quantified the information about all past signals encoded in kinase activity, $\dot{I}_{s\to a}$, and found that *E. coli* use this information efficiently: they climb gradients at speeds near the information-performance limit (1). There are two possible inefficiencies that prevent *E. coli* from reaching the limit: first, cells might encode information about past signals, which don't contribute to gradient-climbing; and second, information about current signal can be lost in communication to the motor behavior. Now that we have an expression for the information about current signal $s(t)$ in kinase activity, we can distinguish between these two effects.

We previously defined the information about all past signals encoded in kinase activity using the following transfer entropy rate:

$$\dot{I}_{s\to a} \equiv \lim_{dt\to 0} \frac{1}{dt} I(a(t+dt); \{s\}|\{a\}) \,. \tag{150}$$

The subset of this information that is relevant to chemotaxis is:

$$\dot{I}^*_{s\to a} \equiv \lim_{dt\to 0} \frac{1}{dt} I(a(t+dt); s(t)|\{a\}) \,, \tag{151}$$

which is the information we have considered here. How do these information rates compare to each other for the kinase response function and noise correlation function that we measured here and previously?

First, note that if kinase activity $a$ were Markovian in $s(t)$, then we would have

$$\begin{aligned} \dot{I}_{s\to a} &= \lim_{dt\to 0} \frac{1}{dt} I(a(t+dt); \{s\}|\{a\}) \\ &= \lim_{dt\to 0} \frac{1}{dt} I(a(t+dt); s(t)|\{a\}) \\ &= \dot{I}^*_{s\to a}, \end{aligned} \tag{152}$$

and all information about signals encoded in kinase activity is relevant to gradient climbing. Surprisingly, this means that a long response adaptation time does not necessarily degrade information about the current signal.

We can evaluate both of these information rates for the response and noise models used here. In the regime of shallow gradients and $\tau_2 \approx \tau_n$, the information about past and present signals is (1,11):

$$\dot{I}_{s\to a} \approx \frac{1}{4\pi} \int_{-\infty}^{\infty} \frac{S(\omega)\frac{r_0^2}{\omega^2}|K_r(\omega)|^2}{N_n(\omega) + r_0\,|K_r(\omega)|^2}\, d\omega \tag{153}$$

$$= \frac{\frac{G_r^2}{\tau_1^2} r_0^2 \, g^2 \, \sigma_v^2 \, \tau_3^2}{4\, D_n \left(1 + \frac{\tau_3}{\tau_v}\sqrt{1 + \frac{G_r^2}{\tau_1^2}\frac{r_0}{2\, D_n}}\right)} = \frac{1}{\tau_v}\frac{1}{4}\frac{R\,\gamma_r \left(\frac{\tau_3}{\tau_v}\right)^2}{\left(1 + \frac{\tau_3}{\tau_v}\sqrt{1+R}\right)} \tag{154}$$

which we have expressed in terms of the ratio of the diffusivity of filtered particle noise and the diffusivity of slow kinase noise, $R = \frac{G_r^2}{\tau_1^2}\frac{r_0}{2\, D_n}$; the particle arrival signal-to-noise ratio, $\gamma_r = 2\, r_0\, g^2\, \sigma_v^2\, \tau_v^3$; and $\tau_3^{-1} = \tau_1^{-1} + \tau_2^{-1}$.

We compare this to the information about current signal only, derived in the previous section, Eqn. 110, reproduced below:

$$\dot{I}_{s\to a}^{*} = \frac{1}{\tau_v}\frac{1}{4}\frac{R}{1+R}\frac{\gamma_r}{\left(1 + \frac{\tau_v}{\tau_3}\frac{1}{\sqrt{1+R}}\right)^2} \tag{155}$$

The ratio of these two information rates has a simple form:

$$\frac{\dot{I}_{s\to a}^{*}}{\dot{I}_{s\to a}} \approx \frac{\tau_v}{\tau_3\sqrt{1+R} + \tau_v} = \frac{\tau_v}{\tau_3\sqrt{1 + \frac{G_r^2}{\tau_1^2}\frac{r_0}{2\, D_n}} + \tau_v}. \tag{156}$$

Thus, for $\dot{I}_{s\to a}$ to mostly carry information about current signal and be close to $\dot{I}_{s\to a}^{*}$, 1) the time scale of initial kinase response must be short compared to the signal correlation time, $\tau_1 \ll \tau_v$; and 2) the diffusivity of filtered particle noise must be small compared to that of internal kinase noise, $G_r^2\, r_0 \ll 2\, D_n$. Using $\tau_1 = 1/60\ s$ from biochemistry studies Refs. (25,26), we estimate that $\frac{\dot{I}_{s\to a}^{*}}{\dot{I}_{s\to a}} \approx 0.88 \pm 0.01$ in $c_0 = 1\ \mu\text{M}$, and increases as $c_0$ gets larger or smaller. This suggests that *E. coli*'s main source of "inefficiency" is that relevant information in kinase activity is lost in communication with the motors.

This result might appear to be in contradiction with the results of Ref. (16), which found that the fraction of predictive information about signals compared to past information about signals was very small (about 1%) in a model of *E. coli*'s kinase activity, $a$, and downstream readout molecules, $x$ (CheYp). Our $\dot{I}_{s\to a}^{*}$, being a predictive information rate, is very similar to their predictive information, while $\dot{I}_{s\to a}$ is very similar to their past information. However, that study considered predictive and past information encoded in the *current* value of the readout molecule, $x(t)$, instead of the entire history of readout molecules $\{x\}$. This difference in how our information quantities are defined explains the large difference in result. Kinase activity $a$ and even CheY phosphorylation level, $x$, are not the final outputs of the chemotaxis system. Downstream pathway dynamics can act on the entire past of $a$ or $x$ to extract information and make behavioral decisions (tumble rate). Therefore, the current values of $a(t)$ and $x(t)$ do not need to be faithful estimates of the current (or future) signal $s(t)$; they just need to carry decodable information about $s(t)$ in their past trajectories. Our information measures above account for this.

#### Summary of information inequalities and results

In summary, we have two sets of inequalities. The first set of inequalities,

$$\dot{I}_{s\to a} \geq \dot{I}^*_{s\to a} \geq \dot{I}^*_{s\to m} \propto \left(\frac{v_d}{v_0}\right)^2, \tag{157}$$

was the focus of our previous work (1), and it quantifies how efficiently *E. coli* use the information *that they have* at the level of kinase activity, $\dot{I}_{s\to a}$, to climb gradients. The main result of that work was that $\dot{I}_{s\to a} \approx 2\,\dot{I}^*_{s\to m}$. The analysis above adds to this: $\dot{I}_{s\to a} \approx \dot{I}^*_{s\to a} \approx 2\,\dot{I}^*_{s\to m}$.

The second set of inequalities,

$$\dot{I}^*_{s\to r} \geq \dot{I}^*_{s\to a} \geq \dot{I}^*_{s\to m} \propto \left(\frac{v_d}{v_0}\right)^2, \tag{158}$$

particularly the left-most one, is the focus of this work. It quantifies how much information *E. coli* get compared to the physical limit. The main result of this manuscript is that $\dot{I}^*_{s\to r} \gg \dot{I}^*_{s\to a}$.

#### Appendix A: Causal Wiener filter derivation

Causal Wiener filtering theory seeks a linear estimator of an unknown quantity $s(t+\tau)$ at time $\tau$ in the future, from past observations of a quantity $x$ that is correlated with $s$ (36). The past of $x$ is denoted $\{x(t)\}$. Both $s$ and $x$ are assumed to be stationary stochastic processes with zero means: $\langle x(t)\rangle = \langle s(t)\rangle = 0$. The Wiener filter, $M_x(T)$, is the kernel that minimizes the mean squared error of the estimator:

$$M_x(T) = \underset{K(T)}{\operatorname{argmin}}\ \langle e^2(\tau)\rangle = \underset{K(T)}{\operatorname{argmin}} \left\langle \left( s(t+\tau) - \int_{-\infty}^{t} K(t-t')\, x(t')\, dt' \right)^2 \right\rangle. \tag{159}$$

In general, the estimator of $s(t+\tau)$ that minimizes the mean squared error is the conditional mean $\langle s(t+\tau)|\{x(t)\}\rangle$. In the case of Gaussian-distributed $s$ and $x$, the conditional mean $\langle s(t+\tau)|\{x(t)\}\rangle$ is exactly a linear function of $\{x(t)\}$, so the linear estimator above is the global optimum. The minimum error $\langle e^*(\tau)^2\rangle = \sigma^2_{s|x}(\tau)$ is the conditional variance of $s(t+\tau)$ given past $x$. The main technical challenge of finding the optimal kernel is the constraint that it must be causal: $M_x(T) = 0$ for $T < 0$.

To derive the optimal kernel, first we expand the square in the objective function:

$$\langle e^2(\tau)\rangle = \left\langle s(t+\tau)^2 - 2\, s(t+\tau) \int_{-\infty}^{\infty} K(t-t')\, x(t')\, dt' + \int_{-\infty}^{\infty} K(t-t')\, x(t')\, dt' \int_{-\infty}^{\infty} K(t-t'')\, x(t'')\, dt'' \right\rangle, \tag{160}$$

and move the expectation inside of the integrals:

$$= \sigma_s^2 - 2\int_{-\infty}^{\infty} K(t-t')\, \langle s(t+\tau)\, x(t')\rangle\, dt' + \int_{-\infty}^{\infty}\int_{-\infty}^{\infty} K(t-t')K(t-t'')\, \langle x(t')\, x(t'')\rangle\, dt'\, dt''. \tag{161}$$

Here we used time-translation invariance of $s$: $\langle s(t+\tau)^2\rangle = \langle s(t)^2\rangle = \sigma_s^2$. Next, change variables to $t' \to \tau' = t - t'$ and $t'' \to \tau'' = t - t''$, replacing absolute time with time delays. $\tau'$ and $\tau'' > 0$ correspond to time delay into the past.

$$= \sigma_s^2 - 2\int_{-\infty}^{\infty} K(\tau')\,\langle s(t+\tau)\,x(t-\tau')\rangle\,d\tau' + \int_{-\infty}^{\infty}\int_{-\infty}^{\infty} K(\tau')K(\tau'')\,\langle x(t-\tau')\,x(t-\tau'')\rangle\,d\tau'\,d\tau''\,. \tag{162}$$

Defining the cross-correlation function $C_{xs}(t-t') = \langle x(t')\,s(t)\rangle$ and autocorrelation function $C_x(t-t') = \langle x(t')\,x(t)\rangle$:

$$= \sigma_s^2 - 2\int_{-\infty}^{\infty} K(\tau')\,C_{xs}(\tau'+\tau)\,d\tau' + \int_{-\infty}^{\infty}\int_{-\infty}^{\infty} K(\tau')K(\tau'')\,C_x(\tau''-\tau')\,d\tau'\,d\tau''\,. \tag{163}$$

When $\tau > 0$ in $C_{xy}(\tau)$, $y$ is evaluated at a time point in the future relative to $x$. Note that $C_{sx}(\tau) = C_{xs}(-\tau)$.

Next, we take the functional derivative of the mean squared error with respect to $K(T)$:

$$\frac{\delta\langle e^2(\tau)\rangle}{\delta K} = -2\,C_{xs}(\tau'+\tau) + \int_{-\infty}^{\infty} K(\tau'')\,C_x(\tau''-\tau')\,d\tau'' + \int_{-\infty}^{\infty} K(\tau'')\,C_x(\tau'-\tau'')\,d\tau'' \tag{164}$$

Since $C_x(\tau) = C_x(-\tau)$, this is:

$$= -2\,C_{xs}(\tau'+\tau) + 2\int_{-\infty}^{\infty} K(\tau'')\,C_x(\tau''-\tau')\,d\tau''\,. \tag{165}$$

Now we need to consider the causal constraint on $K(T)$. For optimality with this constraint, the equation above must equal zero for $\tau' \geq 0$ (at times when $x$ precedes $s$ in $C_{xs}$). Otherwise, for $\tau' < 0$, the derivative is not necessarily zero. Therefore, at the optimum we can write (37):

$$\int_{-\infty}^{\infty} M_x(\tau'')\,C_x(\tau'-\tau'')\,d\tau'' - C_{xs}(\tau'+\tau) = A(\tau') \tag{166}$$

where

$$A(\tau') = \begin{cases} 0, & \tau' \geq 0 \\ a(\tau'), & \tau' < 0 \end{cases} \tag{167}$$

and $a(\tau')$ is some unspecified function. $A(\tau')$ is therefore anti-causal – it is only non-zero at times in the future ($\tau' < 0$). At first glance, this optimality condition might seem less constrained than if $A(\tau')$ were zero for all $\tau'$ (the optimality condition for the optimal non-causal filter). However, the fact that $A(\tau')$ is nonzero for $\tau' < 0$ actually limits the space of filters $M_x(\tau)$ that keep $A(\tau') = 0$ for $\tau' \geq 0$, as we will see below.

Next, we take the Fourier transform of both sides, defined as $f(\omega) = F[f(t)] = \int_{-\infty}^{\infty} f(t)\,e^{i\,\omega\,t}\,dt$. Convolutions in the time domain become element-wise products in the Fourier domain:

$$M_x(\omega)\,C_x(\omega) = C_{xs}(\omega)\,e^{-i\,\omega\,\tau} + A(\omega). \tag{168}$$

On the left-hand side, we have the product of a causal function and a function that is nonzero for positive and negative time delays, the result of which is also nonzero for positive and negative time delays. On the right-hand side, we have a function that is nonzero for positive and negative time delays and an anti-causal function. How do we get the optimal causal kernel $M_x(\omega)$ out of this?

Naively, one might divide both sides by $C_x(\omega)$ and then multiply element-wise by a Heaviside step function in the time domain to get a causal kernel $M_x(\omega)$. However, although the resulting kernel is causal, it does not satisfy the optimality condition. Plugging that kernel back into Eqn. 168, it multiplies the non-causal $C_x(\omega)$, and the result is non-causal. Thus, $A(\omega)$ is non-causal, so that kernel does not satisfy the optimality condition, $A(\tau') = 0$ for $\tau' \geq 0$.

Instead, we need to split $C_x(\omega)$ into causal and anti-causal parts, called a spectral factorization or Wiener-Hopf factorization (17–19):

$$C_x(\omega) = \phi(\omega)\,\phi^*(\omega), \tag{169}$$

where $\phi(\omega)$ is a causal function in the time domain and its complex conjugate $\phi^*(\omega)$ is anti-causal. $\phi(\omega)$ is constructed by putting all poles and zeros of $C_x(\omega)$ with negative real part into $\phi(\omega)$ and those with positive real part into $\phi^*(\omega)$.

Plugging this into the optimality condition:

$$M_x(\omega)\,\phi(\omega)\,\phi^*(\omega) = C_{xs}(\omega)\,e^{-i\,\omega\,\tau} + A(\omega) \tag{170}$$

$$M_x(\omega)\,\phi(\omega) = \frac{C_{xs}(\omega)}{\phi^*(\omega)}\,e^{-i\,\omega\,\tau} + \frac{A(\omega)}{\phi^*(\omega)}. \tag{171}$$

The left-hand side is now a causal function in the time domain, being the product of causal functions, and the right-hand side contains a non-causal function and an anti-causal function.

Multiplying both sides of Eqn. 171 by a Heaviside function in the time domain and then transforming back to Fourier space eliminates the anti-causal term $\frac{A(\omega)}{\phi^*(\omega)}$ and leaves the left-hand side unaffected:

$$M_x(\omega)\,\phi(\omega) = \left[\frac{C_{xs}(\omega)}{\phi^*(\omega)}e^{-i\,\omega\,\tau}\right]^+. \tag{172}$$

Now the right-hand side is causal, and dividing by $\phi(\omega)$ gives the optimal causal filter:

$$M_x(\omega) = \frac{1}{\phi(\omega)}\left[\frac{C_{xs}(\omega)}{\phi^*(\omega)}e^{-i\,\omega\,\tau}\right]^+. \tag{173}$$

To check that this filter satisfies the optimality condition (Eqn. 168), we can plug it in:

$$\frac{1}{\phi(\omega)}\left[\frac{C_{xs}(\omega)}{\phi^*(\omega)}e^{-i\,\omega\,\tau}\right]^+\ \phi(\omega)\,\phi^*(\omega) = C_{xs}(\omega)\,e^{-i\,\omega\,\tau} + A(\omega) \tag{174}$$

$$\left[\frac{C_{xs}(\omega)}{\phi^*(\omega)}e^{-i\,\omega\,\tau}\right]^+\ \phi^*(\omega) = C_{xs}(\omega)\,e^{-i\,\omega\,\tau} + A(\omega) \tag{175}$$

$$\left[\frac{C_{xs}(\omega)}{\phi^*(\omega)}e^{-i\,\omega\,\tau}\right]^+ = \frac{C_{xs}(\omega)}{\phi^*(\omega)}\,e^{-i\,\omega\,\tau} + \frac{A(\omega)}{\phi^*(\omega)}. \tag{176}$$

Now the left-hand side is the causal part of the first term on the right-hand side. Therefore, their difference is anti-causal and $A(\omega)$ is thus anti-causal, as desired:

$$A(\omega) = -\phi^*(\omega) \left[ \frac{C_{xs}(\omega)}{\phi^*(\omega)} e^{-i\,\omega\,\tau} \right]^- . \tag{177}$$

At the optimum, the mean square error $\langle e^2(\tau) \rangle = \sigma^2_{s|x}(\tau)$ is:

$$\sigma^2_{s|x}(\tau) = \sigma^2_s - 2 \int_0^\infty M_x(\tau')\, C_{xs}(\tau' + \tau)\, d\tau' + \int_0^\infty \int_0^\infty M_x(\tau')\, M_x(\tau'')\, C_x(\tau'' - \tau')\, d\tau'\, d\tau'' , \tag{178}$$

where we have set the lower limit to zero because the kernel $M_x(T)$ is zero for $T < 0$. Using the optimality condition $\int_0^\infty M_x(\tau'')\, C_x(\tau'' - \tau')\, d\tau'' - C_{xs}(\tau' + \tau) = 0$ for $\tau' \geq 0$, we get:

$$= \sigma^2_s - \int_0^\infty M_x(\tau')\, C_{xs}(\tau' + \tau)\, d\tau' . \tag{179}$$

Since $C_{xs}(\tau) = C_{sx}(-\tau)$, this is:

$$= \sigma^2_s - \int_{-\infty}^\infty M_x(\tau')\, C_{sx}(-\tau - \tau')\, d\tau' , \tag{180}$$

which is the convolution of $M_x(T)$ and $C_{sx}(T)$, with the result evaluated at $-\tau$. This can be expressed using their Fourier transforms (note the minus sign in front of tau in equation (112) leads to a plus sign in the exponent below) as:

$$\sigma^2_{s|x}(\tau) = \sigma^2_s - \frac{1}{2\pi} \int_{-\infty}^\infty M_x(\omega)\, C_{sx}(\omega)\, e^{i\,\omega\,\tau}\, d\omega . \tag{181}$$

Plugging in the optimal kernel:

$$= \sigma^2_s - \frac{1}{2\pi} \int_{-\infty}^\infty \frac{C_{sx}(\omega)}{\phi(\omega)} \left[ \frac{C_{xs}(\omega)}{\phi^*(\omega)} e^{-i\,\omega\,\tau} \right]^+ e^{i\,\omega\,\tau}\, d\omega . \tag{182}$$

Finally, the correlation coefficient is:

$$\rho^2_{xs}(\tau) = 1 - \frac{\sigma^2_{s|x}(\tau)}{\sigma^2_s} = \frac{1}{\sigma^2_s} \frac{1}{2\pi} \int_{-\infty}^\infty \frac{C_{sx}(\omega)}{\phi(\omega)} \left[ \frac{C_{xs}(\omega)}{\phi^*(\omega)} e^{-i\,\omega\,\tau} \right]^+ e^{i\,\omega\,\tau}\, d\omega . \tag{183}$$

#### Supplementary Figures

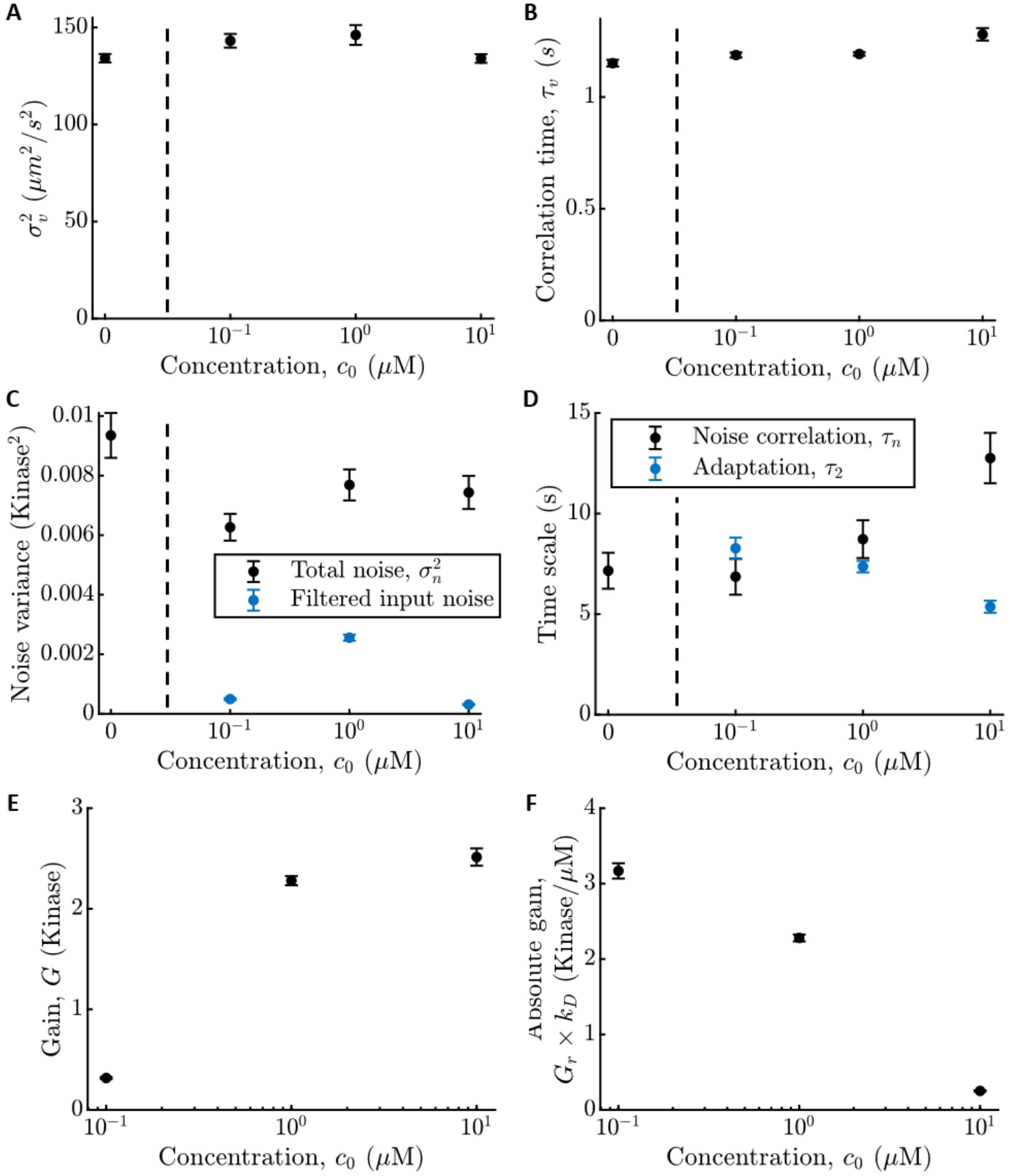

**Figure S1: Measured signal, response, and noise parameter values in different background concentrations. A)** Estimated variance of up-gradient velocity, $\sigma_v^2$, as a function of background $c_0$, which together with the gradient steepness $g$ sets the signal strength. Horizontal axes are on log-scale, and vertical dashed lines throughout separate parameters measured at $c_0 = 0$ from those measured at finite

$c_0$. Error bars throughout are standard error of the mean (see Methods section of the main text). **B)** Correlation time of up-gradient velocity, $\tau_v$, which sets the signal correlation time. Parameters in (A) and (B) are those of the median phenotype in Fig. S2, with tumble bias $TB \approx 0.09$. **C)** Variance of the total noise in kinase activity, $\sigma_n^2$ (black), and the estimated variance of particle arrival noise filtered through the kinase response kernel (blue) with $\tau_1 = 1/60\ s$ (25,26). **D)** Kinase noise correlation time, $\tau_n$, and kinase response adaptation time, $\tau_2$ (blue). **E)** Gain of kinase response to signal or log-concentration, $G$. **D)** Gain of kinase response to absolute concentration, $G_c = k_D\, G_r = G/c_0$, where $G_r$ is the gain of kinase responses to particle arrival rate.

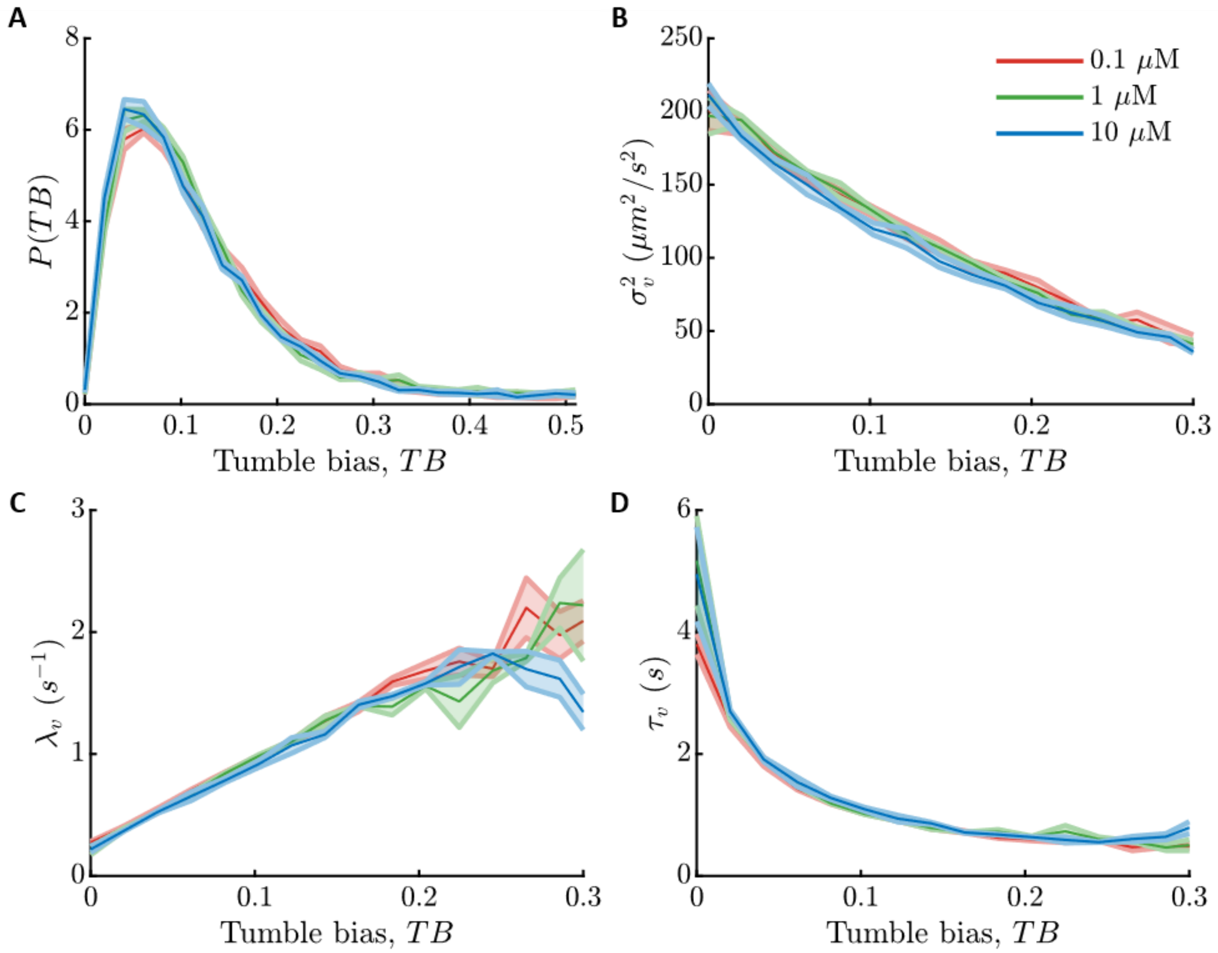

**Figure S2: Swimming parameters as a function of tumble bias in different background concentrations. A)** Distribution of tumble bias, $TB = 1 - P_{run}$, or fraction of time cells spend in the tumble state, among cells in an isogenic population. Throughout: red is $c_0 = 0.1$ μM, green is $c_0 = 1$ μM, and blue is $c_0 = 10$ μM. Shading is standard error of the mean (Methods). **B)** Variance of up-gradient velocity, $\sigma_v^2$, versus tumble bias, $TB$. **C)** Velocity decorrelation rate, $\lambda_v = \tau_v^{-1} \approx (1 - \alpha)\, \lambda_{R0} + 2\, D_r$, versus $TB$. $\alpha$ quantifies how correlated heading is before and after a tumble; $\lambda_{R0}$ is the average tumble rate; and $D_r$ is the rotational diffusion coefficient (1). **D)** Velocity correlation time, $\tau_v$.

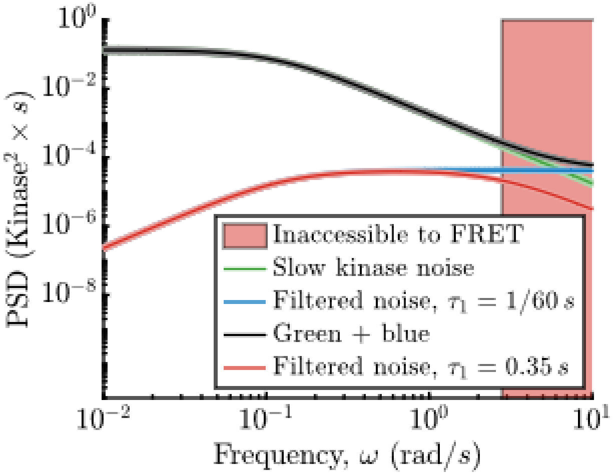

**Figure S3: Noise power spectra.** In frequency space, kinase responses to particle arrivals implies that the noise in kinase activity must be larger than filtered particle arrival noise (blue, using $\tau_1 = 1/60\ s$ from biochemistry studies Refs. (25,26)). At low frequencies where we can measure noise and responses with our FRET system (green), this bound is far from saturated. Naively extrapolating to higher frequencies (red shaded region, marked by the value of $1/\tau_1$ measured in FRET experiments) violates this bound (the green line goes below the blue line). This implies either additional noise at high frequencies that is not captured by a single exponential (black line is slow noise, green, plus filtered particle noise, blue) or a slower kinase response time $\tau_1$ (red line is filtered particle noise with $\tau_1 \approx 0.35\ s$ measured in FRET experiments), which could be a necessary by product of the coupling between kinases that creates large gain (thus raising the red line) but also slows down the response. The behaviorally-relevant information rates computed in the main text are insensitive to these choices.

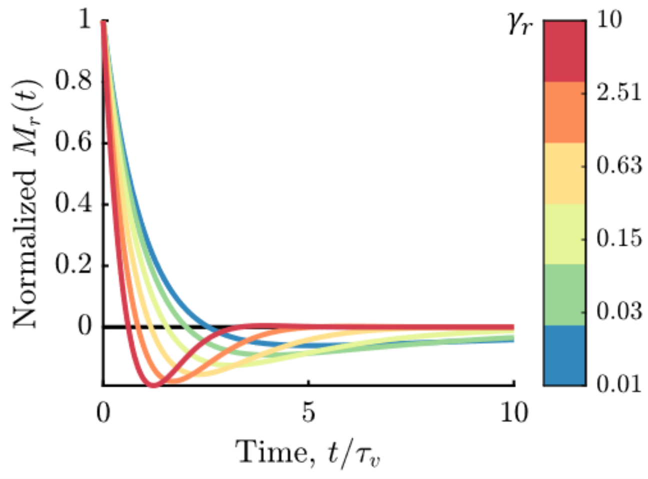

**Figure S4: Optimal kernel for inferring current signal, $s(t)$, from past particle arrivals, $r$.** Colors indicate different values of the signal-to-noise ratio $\gamma_r$, marked on the right. Each kernel is normalized so that $M_r(0) = 1$.

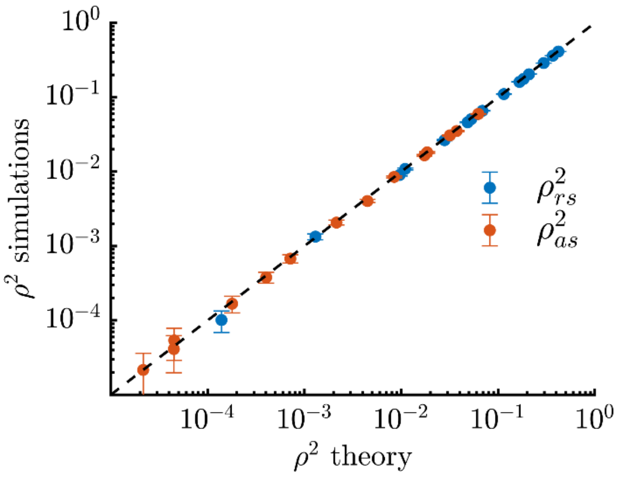

**Figure S5: Squared Pearson correlation coefficients, $\rho^2_{rs}$ and $\rho^2_{as}$, between true signals $s(t)$ and optimal estimates, $\hat{s}_r(t)$ and $\hat{s}_a(t)$, computed from simulations and theory.** Dots are varying $c_0 \in \{0.1, 1, 10\}$ µM and $g \in \{0.01, 0.1, 0.2, 0.3, 0.4\}$ $\mathrm{mm}^{-1}$. Simulations are described in SI section **Simulation details**. Values of the squared correlation coefficients range from $10^{-4.6}$ to $0.42$. Error bars are SEMs and were computed by bootstrapping trajectories. The theoretical expressions show excellent agreement with the correlations computed from simulations.
